# Invivo and systematic analysis of random multigenic deletions associated with human diseases during epithelial morphogenesis in Drosophila

**DOI:** 10.1101/2021.01.20.427453

**Authors:** Usha Nagarajan, Shanmugasundaram Pakkiriswami, Sandiya Srinivasan, Niveditha Ramkumar, Rajeswari Rajaraman, Kumarasamy Thangaraj

**Author notes:** Equal contribution (Usha Nagarajan, Shanmugasundaram Pakkiriswami).

## Abstract

Random loss of multigenic loci on chromosomes, a crucial drive for evolution, occurs frequently in all living organisms. Analysis of such chromosomal disruption and understanding the consequences of their impact on the growth and development of multicellular organisms is challenging. In this report, we have addressed this issue using invivo mosaic analysis of deficiency lines in Drosophila. Genes on fly deficiency lines were compared with human orthologs for their implications in disease development during cytoskeletal processes and epithelial morphogenesis. The cytoskeletal phenotypes from the fly has been utilized to predict the function of human orthologs. In addition, as these Drosophila deficiency lines are equivalent to human microdeletions, based on the clonal behaviour and phenotypes generated, a systematic analysis has been carried out to establish the critical loci that correspond to Microdeletion Syndromes and Mendelian Disorders in humans. Further we have drawn the synteny that exists between these chromosomes and have identified critical region corresponding to defects. A few potential candidates that might have an implication in epithelial morphogenesis are also identified.

## Introduction

Contiguous-gene deletion syndromes or multigenic deletion syndromes display extremely variable clinical symptoms depending on the genes lost in the loci (Rauch et al., 2001; Slavotinek, 2012). A great deal of evidence shows that random loss of genes or spontaneous changes on chromosomes (like virus-mediated mutagenesis in humans causing chromatid breakages, translocations, duplications, deficiencies/deletions leads to developmental anomalies and growth defects (Fortunato and Spector, 2003). “Less-is-more” hypothesis insists that gene loss is an engine for evolutionary change and disease development (Olson, 1999; Olson and Varki, 2003). Despite the fact that the adaptive role of gene loss has formed the basis for evolution, impact of their adaptive biological functions on individual integrity and stability remains elusive. Just as how loss and gain of Homeotic genes and their clusters prove fundamental for the evolution of body shape and size (Ruddle et al., 1994), the loss and gain of genes implicated in epithelial morphogenesis also influence the integrity of body size and shape (Schock and Perrimon, 2002). In a multicellular organism, epithelial morphogenesis is an inevitable process in which epithelia, tissues that line the organ and body surfaces, determine organ-to-body shape and size (<u>Gumbiner BM</u>.1992; Schock and Perrimon, 2002). Of all the factors implicated in this process, cytoskeletal components that balance rigidity and flexibility of the cell, play a critical role in maintaining organ size and growth of the organism (Knust, 2005; Gibson and Perrimon, 2005; Shen and Dahmann, 2005; Donohoe et al., 2018). Defects during this process lead to loss of tissue integrity; development of genetic disorders and diseases (Olson and Varki, 2003; Knust, 2005 in Birchmeier and Birchmeier, 2005).

Compared to loss of individual essential genes and frameshift mutations, gains and losses of multiple (more than one) genes bring about greater dramatic genetic repercussions. Random loss of contiguous (loss of genes in the same order) genes would disrupt the normal development, affecting growth and patterning of the tissue, which otherwise occur concurrently. Loss of such contiguous genes that may occur due to any kind of genetic aberrations might regulate both cell-autonomous and non-autonomous functions of the tissue giving rise to heterogeneity in cell population (mosaicism). It is intriguing to understand the influence of such sudden, random loss of genes causing mosaicism and to compare their effect with other organisms.

Genomic studies suggest that genetic mosaicism is a very common phenomenon during epithelial morphogenesis. Although such lesions to the genome are random and generally asymptomatic in initial stages, affected individuals subsequently develop varying levels of metabolic disorders and disabilities (Chial, 2008). Also, since such incidences occur sporadically, chances of undertaking systematic analysis is very low or may not be feasible. In the past, rigorous systematic analysis of genes (Reiter et al., 2001) and genetic screening (St Johnson, 2002; Yamamoto et al., 2014) have been undertaken to understand genes implicated in disease development and their functions. However, how these individuals subjected to such sudden and random loss of genes manage without their gene function and what impact does it have on the individual is not clear. In multigenic deletions, the synergistic and antagonistic effect of genes in maintaining homeostasis or disease development and the alternate pathways that operate to compensate the loss is not known yet. Although it is intriguing to understand the consequence of random loss of multiple genes that leads to defects and genetic disorders, it is extremely challenging.

We hypothesized that those limitations can be overcome and unanswered questions can be addressed by using the available *Drosophila* deficiency line collections, one of the unique genetic tools well-utilized extensively for genetic screening, mapping and characterization processes. Two types of molecularly defined deficiency lines, whose details including ends of the genes lost as well as cytological location on the chromosome are known, are available (Ryder et al., 2004; Parks et al., 2004; Cook et al., 2012; Roote and Russell, 2012). In addition, as chromosomes of these deletion-bearing fly stocks resemble those chromosomes with microdeletion or chromosomal deletions of contiguous genes (loss of genes in the same order) in other organisms, the collection has been utilized for systematic and comparative analysis to establish the critical loci that correspond to Mendelian Disorders and Microdeletion Syndromes in humans.

Based on this logic, in this work, we set out a pilot project by subjecting these deficiency lines for mitotic clonal analyses in *Drosophila* wing imaginal discs to identify and understand their role in cytoskeletal organization during epithelial morphogenesis. Clonal analysis technique allows to analyse behaviour of two different population of cells in a tissue (Lee and Luo, 2001; Wu and Luo, 2006; Smith-Bolton et al., 2009; Rodriguez et al., 2012; Powell and Piddini, 2016). Here we have utilized this property to understand the behaviour of randomly eliminated genes, thereby identify the critical gene in that locus implicated in epithelial morphogenesis. During this process of clonal analysis, we have also followed the cytoskeletal defects produced by these deficiency lines. We also undertook a systematic functional analysis of the genes in the fly deficiency lines, compared them with human orthologs and analysed their implications in cytoskeletal processes and epithelial morphogenesis. Further we have drawn the synteny that exist between these chromosomes and identified critical region of epithelial defects. Here, in this report, few potential candidates that might have an implication in epithelial morphogenesis are discussed.

## Results and Discussion

### (i) Recombination of Deletion-bearing chromosomes with FRT chromosomes and generation of mitotic clones

To address the behavioural property of the tissue subjected to multigenic deletions we generated mitotic clones using the available deficiency lines in Drosophila wing imaginal discs. We recombined X and II chromosome deficiency lines with their respective FRT chromosomes. We have subjected ∼67 FRT-recombined X chromosome deficiency lines and 125 FRT-recombined second (2L and 2R) chromosome deficiency lines for clonal analysis in III instar larval wing imaginal discs. These deficiency lines produced clones of varied number and sizes (Figure 1(A-O); Supplementary S1, Sheet2 – Sheet5). For instance, deficiency line Df(1)Ex6237(BL-7711) with 21 genes deleted produced 103 clones, highest average number of clones (Figure 1N) while deficiency line Df(1)ED6443(BL-9053) with 45 genes deleted generated 0 to 1 (average =0.3), lowest number of clones. Deficiency line Df(1)Exel6230(BL-7705) with 9 genes deleted developed the biggest clones of 0.6 Sq. pixels, while deficiency line Df(1)Exel6221(BL-7699) with 14 genes deleted produced smallest clones of 0.04 Sq. pixels (Figure 1J), Supplementary S1-Sheet2-3). Some of the deficiency lines did not generate clones or generated few clones (<10) per wing disc. Interestingly few of the deficiency lines showed clones specifically localized in wing discs, especially only in pouch region (Figure1J) or only in notum of the wing disc (Figure 1F, Supplementary 2(A-B).

**Figure 1.**
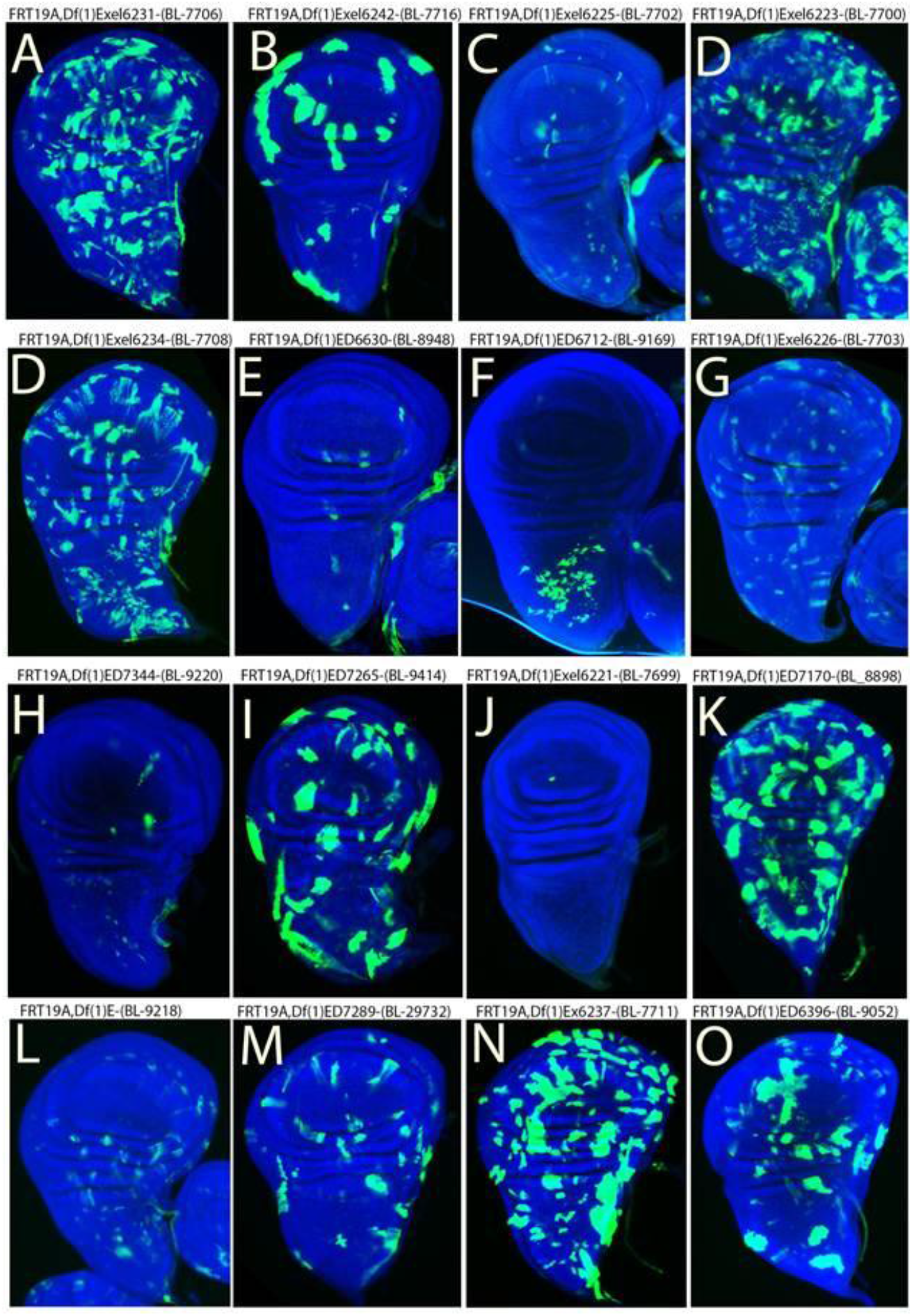
Distribution and size of MARCM clones. Representative Wing discs (A-O) with Mitotic clones (GFP positive, counterstained with DAPI in blue) generated using Deficiency lines on X chromosome recombined to FRT19A chromosomes. Note the variation in size and distribution of the clones in wingdiscs.

In order to understand the clonal properties of the mitotic clones generated by these deficiency lines, we have categorized the deficiency lines into very few (rare or less than or upto 10 number of clones), few (upto an average of 10-30 clones), moderate (upto 60 clones), and numerous (> 60 clones) clonal types and based on the size of clones, deficiency lines were grouped into large, medium and small types. On X chromosome, 37% of the deficiency lines generated rare/very few clones and 27% generated few clones. Around 21% and 15% of deficiency lines were categorized into moderate and numerous categories respectively (Figure 2A, Supplementary S1-Sheet2). In addition, based on the average size of the clones generated per wing disc, we have categorized the deficiency lines from small through medium to large (Figure 2B, 2(E-F), Supplementary Data S1 Sheet3).

**Figure 2.**
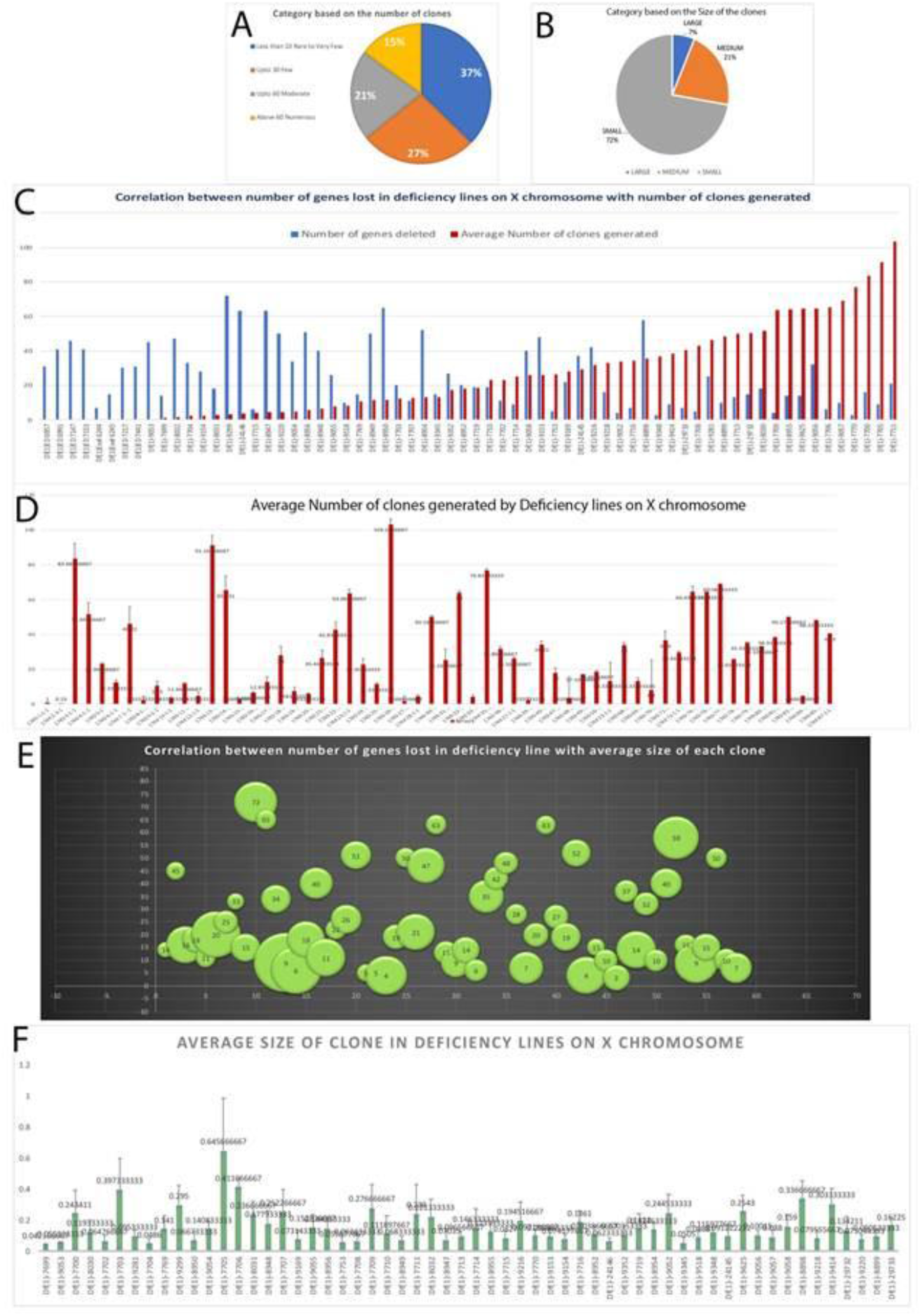
Number and size of clones generated by deficiency lines on X chromosome. Classification of clones based on number of clones (A) and size of the clones (B). Note average number (D) and size of mitotic clones (F) generated by deficiency lines spanning the X chromosome. Correlation between number of genes lost in deficiency lines on X chromosomes with average number (C) and size of mitotic clones (E) generated by them.

To address the link between the number of genes lost in a deficiency line and their corresponding clonal property, we have generated correlation graphs between them. From the correlation graph, we find that depending on the function and importance of genes deleted in a deficiency line, their clonal properties and behaviour varied accordingly (Figure 2C, Supplementary S1-Sheet2). For example, (i) on X chromosome there are four deficiency lines that have very less number of genes lost on them but generated either rare clones [Deficiency line (Df(1)Exel6244(BL-7717) which lost only 7 genes did not generate clones at all while Df(1)Exel6221(BL-7699)which lost 14 genes generated very less to 1 clone in a wing disc (Figure1J)] or very few clones [Df(1)Ex6241(BL-7715) lost 6 genes but generated on an average of only 4 clones (Supplementary Figure2C) while Df(1)ED6579(BL-9518)(70.1) lost 10 genes but generated on an average of 10 clones per wingdisc (Supplementary Figure2D)]. (ii) Similarly, there are only four deficiency lines in the moderate and numerous categories with more than 20 genes lost in them. Deficiency line Df(1)ED7170(BL-8898) lost 58 genes made on an average 35 clones per wingdisc(Figure1K) and Df(1)ED6521(BL-9281) lost 25 genes generated on an average 46 clones per wingdisc (Figure(1L). Similarly, in the numerous category, Df(1)ExelED6989(BL-9056)(76-1) that lost 32 genes made on average of 65 clones per wingdisc (Supplementary Figure 2G). Of all these, notable ones are Df(1)Exel6237(BL-7711) that lost 21 genes that made 103 clones, the highest average number of clones per wingdisc (Figure1N) and Df(1)Exel6223(BL-7770) that had lost only 3 genes but made 77 clones on an average in a wingdisc (Supplementary Figure 2H). This deficiency line had the least number of genes lost with the fourth highest number of clones (Supplementary S1-Sheet2). There are nearly 8 deficiency lines on the X chromosome and 26 deficiency lines on the II chromosome that did not generated clones (Figure 2A and 2C, Figure 3A Supplementary S1 Sheet 2 and 4).

**Figure 3.**
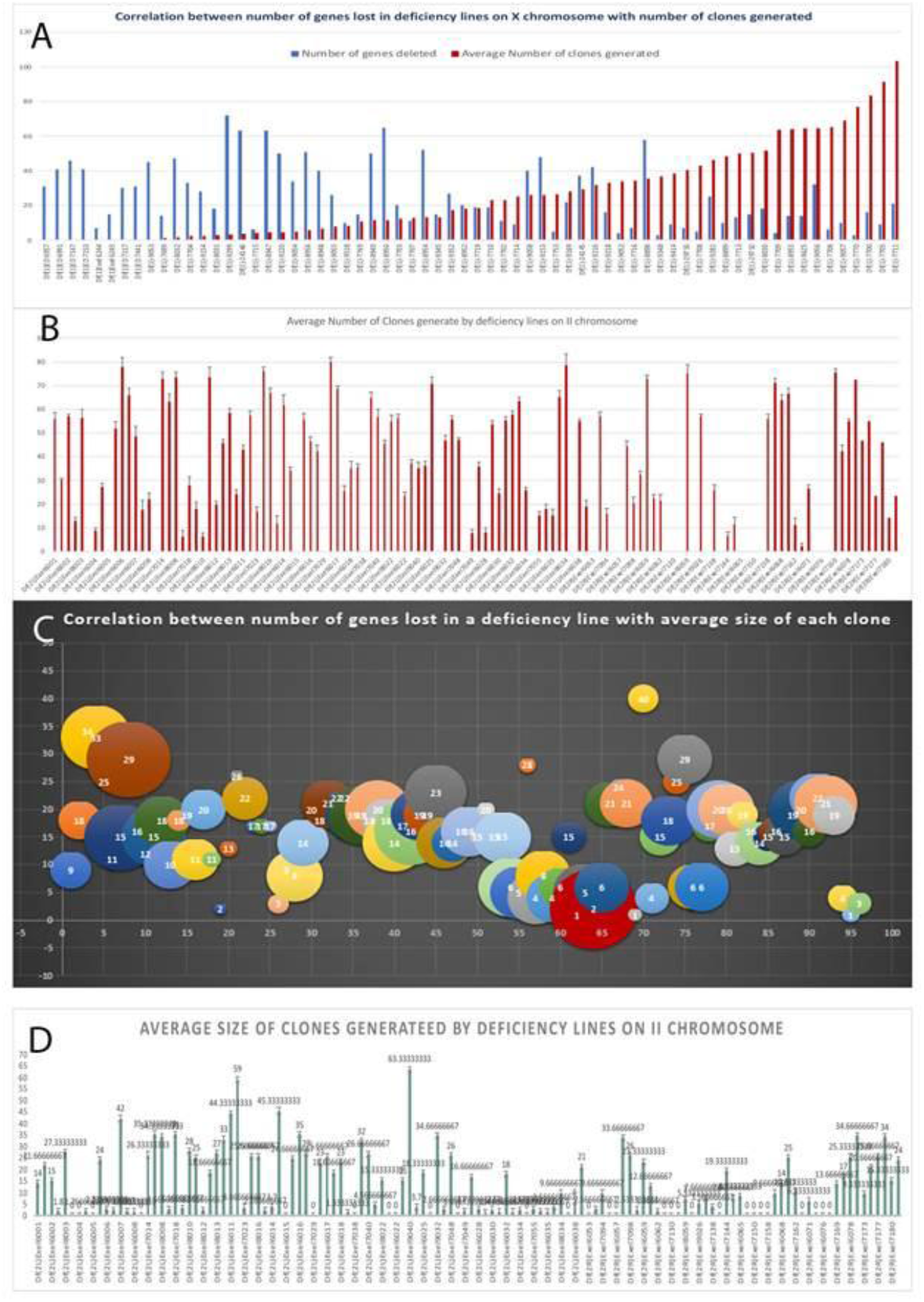
Number and size of clones generated by deficiency lines on II chromosome. Correlation between number of genes lost in deficiency lines on II chromosomes with average number (A) and size of mitotic clones (C) generated by them. Note average number (B) and size of mitotic clones (D) generated by deficiency lines spanning the X chromosome.

Similarly, we have performed clonal analysis on deficiency lines on II chromosome and categorized them based on their number, size and distribution (Figure 3(A-D) Supplementary Data S1 Sheet4-5). Nearly 27% of the deficiency lines were in rare/ very few category while 23% of them were in few category. Around 31% and 19% of deficiency lines were categorized into moderate and numerous categories respectively. As in case of the X chromosome, the number of clones generated by corresponding deficiency line depended on the gene type and function. The deficiency line Df(2L)Exel6024, that is deficient in 19 genes, makes the smallest clones that have an average size of 0.6 Sq. pixels. The largest clones, with a mean size of 63, are made by the Deficiency line Df(2L)Exel9040, that lacks only 2 genes. The line Df(2L)7042, that has lost 14 genes, makes on average, clones sized 15 Sq. pixels while the line Df(2R)Exel6065, that has lost about 40 genes, makes but relatively small clones of average size 8 Sq. pixels (Figure 3(A-D) Supplementary Data S1 Sheet4-5). These findings reinforce only the hypothesis further that the clonal phenotype exhibited by the animals under study depends on the function of the genes lost and their importance to the development underway. We have utilized these clonal properties for systematic and comparative analyses discussed in next sections to understand their implications on human diseases.

### 2. Analysis of cytoskeletal changes associated with Drosophila deficiency lines and functional analysis of their Human orthologs

To understand the role of genes in deficiency lines during epithelial morphogenesis, we have analysed the cytoskeletal changes mediated by generating mitotic clones in wing imaginal discs. Of the 66 deficiency lines on X chromosomes subjected to clonal analysis, nearly 12 deficiency lines showed cytoskeletal changes in the wing imaginal discs (Table 1). Although we generated clones for both X and II chromosomes, we analysed the cytoskeletal defects for entire X chromosomes deficiencies and part of the II chromosome deficiency lines that showed visible defects either in larval or adult stages.

**Table 1:**
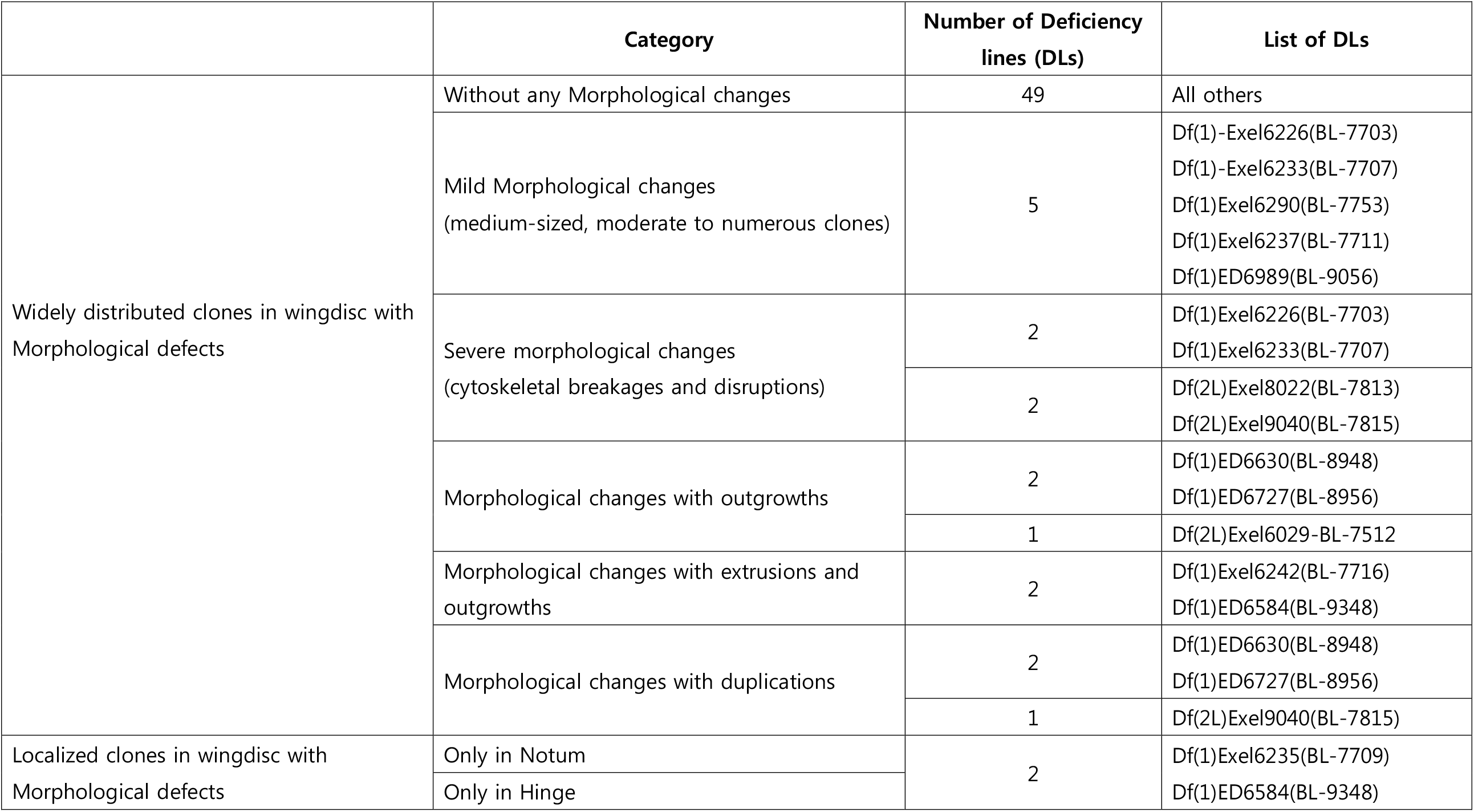
Category based on Morphological/cytoskeletal changes

Based on the type and the degree of cytoskeletal changes, we found that clones were either widely-distributed throughout the wingdisc or localized in wingdiscs. Accordingly, we have classified them into six categories, namely, with (i)-no, (ii)-mild, (iii)-severe morphological changes and with (iv)-outgrowths, (v)-extrusions and (vi)-duplications (Table 1). Nearly 49 deficiency lines did not show any cytoskeletal changes. Of the remaining deficiency lines, approximately 5 deficiency lines (Df(1)-Exel6226(BL-7703), Df(1)-Exel6233(BL-7707), Df(1)Exel6290(BL-7753), Df(1)Exel6237(BL-7711), Df(1)ED6989(BL-9056) showed very mild morphological changes. These deficiency lines generated medium-sized, moderate to numerous clones in the wing discs. In the Df(1)Exel6226(BL-7703), we could observe cell-autonomous cytoskeletal defects (Figure 4B(1-4)) while the deficiency line Df(1)Exel6233(BL-7707), displayed non-cell autonomous cytoskeletal defects involving cytoskeletal breakages and disruptions (Figure 4C(1-4)). Similarly, for second chromosomes, we generated clones using conventional method. Compared to the control wingdiscs (Figure 4D(1-4)), two deficiency lines Df(2L)Exel8022(BL-7813) (Figure 4E(1-4)) and Df(2L)9040(BL-7815) (Figure 4F(1-4)) showed severe cytoskeletal defects. Two deficiency lines (Df(1)Exel6242(BL-7716) Figure 5(A’-A”, Figure 5C’))and Df(1)ED6584(BL-9348) (Figure 5(B’-B”, Figure 5C”) showed cytoskeletal defects and extrusion of the clones associated with outgrowths. Consistently, we find both deficiency lines show similar phenotypes by both direct mosaic as well as MARCM techniques. Two deficiency lines showed localized clones: Df(1)ED6584(BL-9348) produced clones near the periphery of pouch region associated with the extrusion of rounded clonal cells (Figure 5B” and 5C”); while the deficiency line Df(1)Exel6235(BL-7709) produced clones only in the notum region displaying non-cell autonomous cytoskeletal rounding near the clones, (Figure 5(D’-D”)). Similarly, three deficiency lines showed cytoskeletal defects with outgrowths and duplication phenotypes. Of them, two from X chromosome, (Df(1)ED6630(BL-8948) (Figure 6(B-D) and Df(1)ED6727(BL-8956) (Figure 6(E-F) rarely produced clones, however, it showed severe morphological changes, including outgrowth and duplication in the wingdiscs. One of the II chromosome deficiency lines, Df(2L)9040(BL-7815), displayed outgrowth and pouch duplication phenotype in the wingdisc (Figure7(A-D).

**Figure 4.**
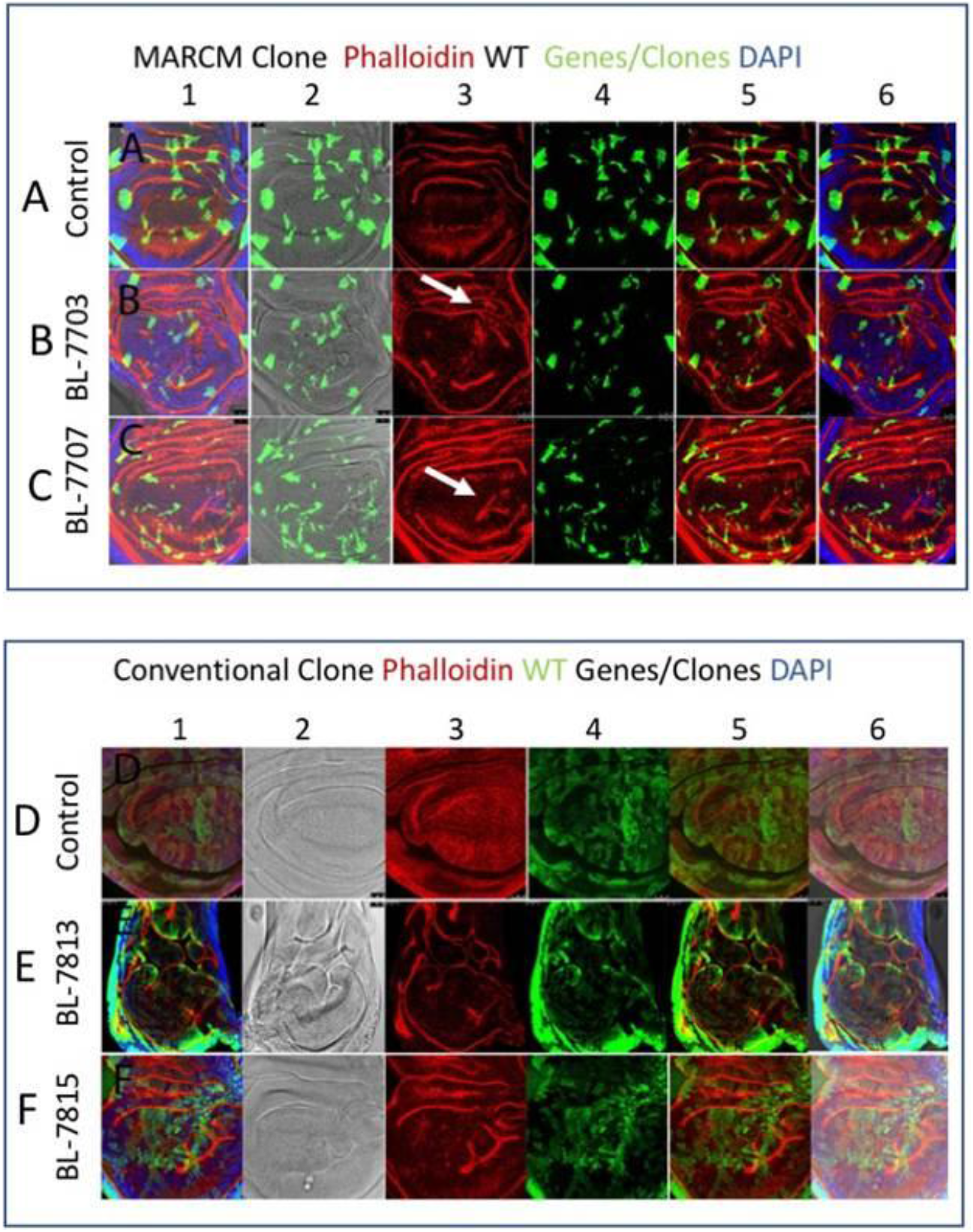
Wing discs with cytoskeletal breakages and disruptions. Compared to control wingdiscs with MARCM clones (A1-A6), GFP positively marked clones by X chromosome deficiency line BL-7703 display cell autonomous (B1-B6) and deficiency line BL-7707 display non-cell autonomous (C1-C6) cytoskeletal defects as shown by Actin (Phalloidin, Red). Compared to control wingdiscs with conventional clones (D1-D6), GFP negatively marked clones by II chromosome deficiency line BL-7813 display cell autonomous (E1-E6) and deficiency line BL-7815 display non-cell autonomous (F1-F6) cytoskeletal defects as shown by Actin (Phalloidin, Red). All panels are standard confocal XY sections. Note panels B3, C3, E3 and F3 for the changes in Actin (Red) representing cytoskeletal breakages and disruptions in wingpouch.

**Figure 5.**
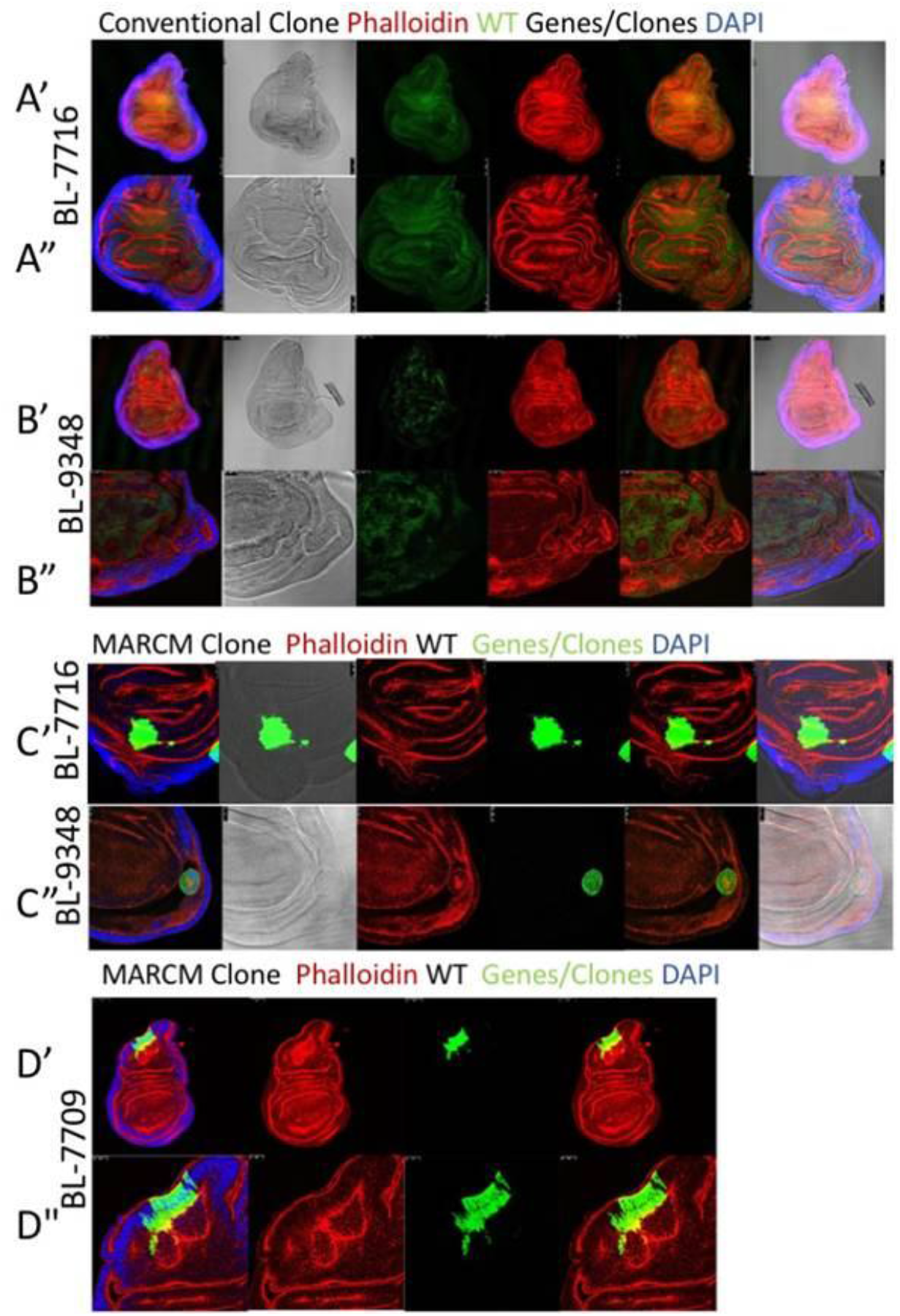
Wing discs with cytoskeletal defects and extrusions. Two deficiency lines BL-7716 (A’-A”, 5C’) and BL-9348 (B’-B”, 5C”) show similar and consistent cytoskeletal defects in clones generated either by conventional method or MARCM strategies. Cells and tissue adjacent to the clones are extruded out of the surface. Localized cytoskeletal defects are produced either along the periphery of the pouch region as shown by deficiency lines BL-9348 (5B” and 5C”) or only in the hinge region as shown by deficiency line BL-7709(D’-D”). Note the rounding of the clones and associated with developmental defects.

**Figure 6.**
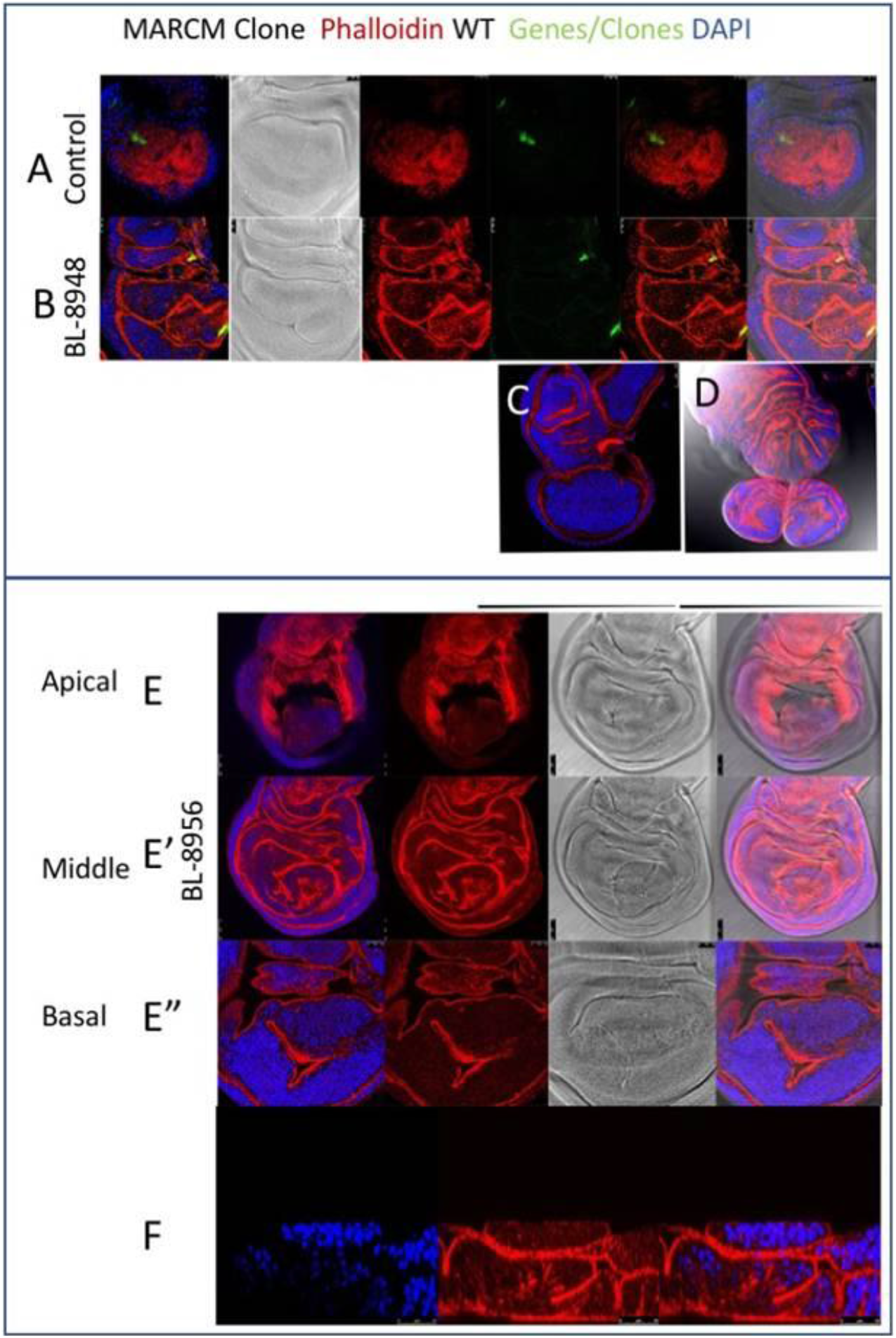
Wing discs with severe cytoskeletal defects associated with outgrowths and duplications. Compared to the control (A) rare clone produced by two deficiency lines BL-8948 (B) and BL-8956 (E-E”). Note the outgrowth of the wingpouch in apical (E), medial (E’) and basal (E”) XY sections and XZ section of the medial region (F).

We have also undertaken a systematic functional analysis of the genes in the fly deficiency lines, compared them with human orthologs and analysed their implications in cytoskeletal processes and epithelial morphogenesis (Supplementary Data S2). In the X chromosome of the fly, out of 1103 total genes, there are 181 genes that are involved in such functions. Among the 181 genes, one-third includes transmembrane proteins, another one-third is involved in cytoskeleton binding and organization, and others are involved in intracellular vesicular transport, cell polarity establishment and maintenance of epithelial morphogenesis. Similarly, in the II chromosome there are 431 genes out of 1245 which are implicated in cytoskeletal functions. We have compared these genes on X and II chromosomes that have cytoskeletal implication with their respective Human orthologs (Figure 8). In humans, 200 of the total orthologs are involved in cytoskeletal functions. In some instances, the functions of the orthologs of cytoskeletally implicated genes are known while in others it is known atleast in the fly counterparts. About 122 genes and their orthologs both have cytoskeletal functions. In numerous cases, the human orthologs of such genes have not been functionally characterized as yet. For 18 genes on X chromosome, cytoskeletal functions of fly orthologs are known but unknown in human orthologs (Table 2).

**Table 2:** Comparison of fly genes implicated in cytoskeletal functions with corresponding human orthologs with unknown function

| <u>DEFICIENCY LINE</u> | <u>FLY GENE</u> | <u>HUMAN GENE</u> | <u>OMIM</u> | <u>CYTOSKELETAL IMPLICATION</u> |  |
| --- | --- | --- | --- | --- | --- |
|  |  |  |  | <u>IN FLY</u> | <u>IN HUMANS</u> |
| Df(1)ED6443 | SKPA | SKP1 | 601434 | Microtubule Cytoskeletal Organization | – |
| Df(1)ED6330 | kirre | KIRREL3; kirre like nephrin family adhesion molecule 3 | 607761 | Cell Adhesion, Plasma Membrane Fusion | NIL |
| Df(1)Exel6227 | CG11448 | RILP1 | 614092 | Dynein Light Chain And Small GTPase Binding |  |
| Df(1)Exel8196 | inc | KCTD5 | 611285 | Cytoskeletal Binding | NIL |
| Df(1)ED6565-9299 | pcx | PCNX1; pecanex homolog 1 | 617655 | Notch Pathway | Transmembrane Protein |
| Df(1)Exel6233BL7707, Df(1)ED6712-BL9169 | CG10804 | TROVE2; TROVE domain family member 2 | 600063 | Transporter | NIL |
| Df(1)Exel6233BL7707, Df(1)ED6712-BL9169 | CG16781 | MARCH5; membrane associated ring-CH-type finger 5 | 610637 | Nil | NIL |
| Df(1)Exel6290 | NAAT1 | SLC6A5; solute carrier family 6 member 5 | 604159 | Sodium Amino Acid Symporter | NIL |
| Df(1)ED7067 | Drak | STK17B; serine/threonine kinase 17b | 604727 | Epithelial Morphogenesis, Stimulates Actomyosin Contractility | NIL |
| Df(1)ED7217 | CG32628 | AGBL2; ATP/GTP binding protein like 2 | 617345 | Mitochondrial Organization | NIL |
| Df(1)ED7217 | NnaD | AGBL2; ATP/GTP binding protein like 2 | 617345 | Mitochondrial Organization | NIL |
| Df(1)ED6716 | brn | B3GNT3 | 605863 | Border Follicle Cell Migration, Maintenance | NIL |
|  |  |  |  | Of Epithelial Cell Polarity, Cell Adhesion And Morphogenesis |  |
| Df(1)ED7165 | CG12717 | SENP7 | 612846 | Axonogenesis,Dendritic Spine Morphogenesis | NIL |
| Df(1)ED7165 | Bap60 | SMARCD1 | 601745 | Myosin Binding | NIL |
| Df(1)ED7170-BL8898 | Sno | SBNO1 | 614274 | Notch Regulating | NIL |
| Df(1)ED7261 | CG5334 | MKRN1; makorin ring finger protein 1 | 607754 | GTPase, Vesicle Fusion Regulation, Intracellular Protein Transport | NIL |
| Df(1)ED7261 | CG32595 | HARBI1; harbinger transposase derived 1 | NIL | Centripetal Follicle Cell Migration | NIL |
| Df(1)ED7344 | CG8509 | PTPA; protein phosphatase 2 phosphatase activator | 600756 | Mitotic Spindle Organization | NIL |
| Df(1)ED7344-9220 | Efhc1.1 | EFHC2; EF-hand domain containing 2 | 300817 | Synaptic Growth, Dendritic Morphogenesis | NIL |

**Figure 7.**
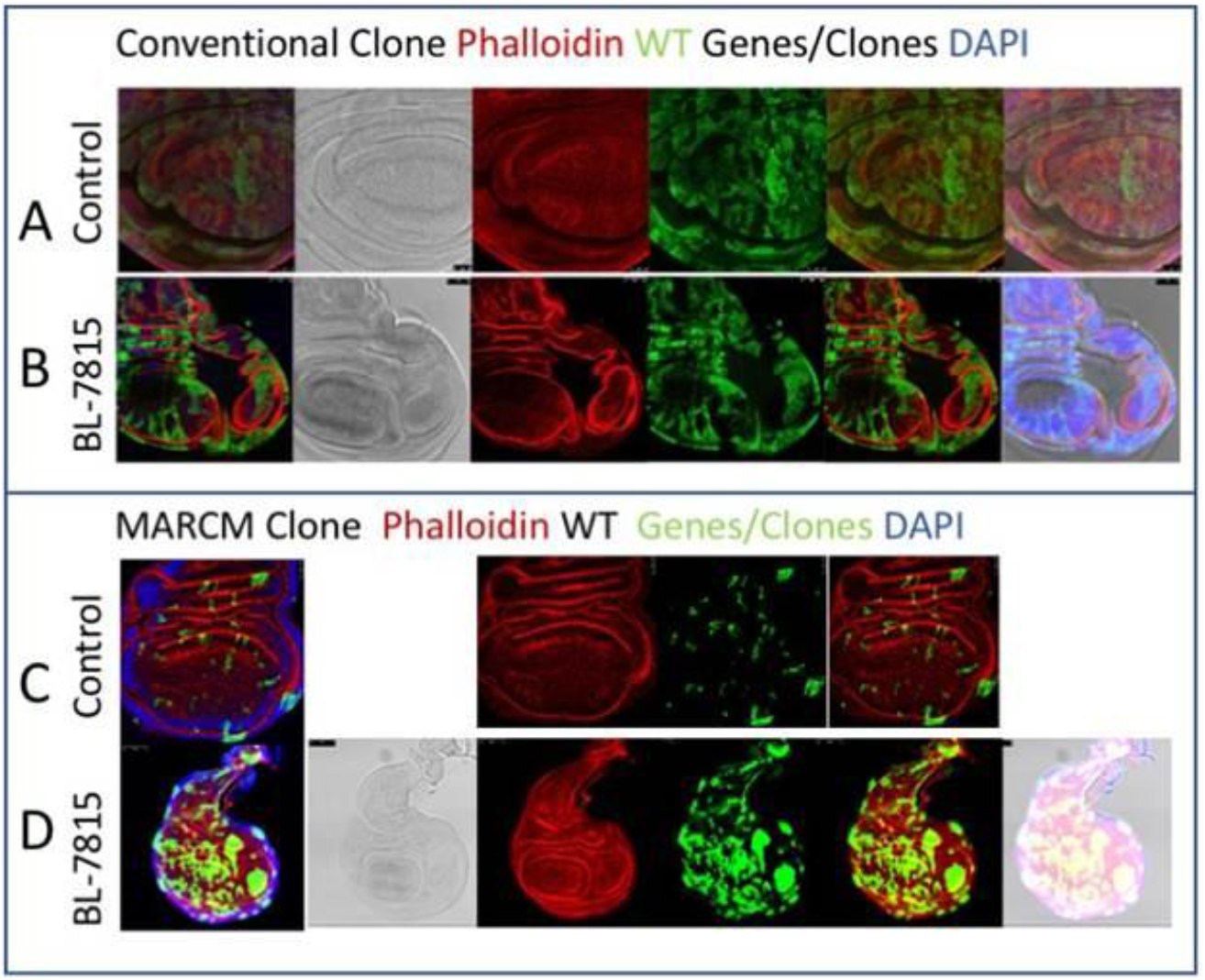
Wing discs with severe cytoskeletal defects associated with outgrowths and duplications. Compared to control wingdiscs with conventional clones (A) or MARCM clones(C) clones generated by deficiency line BL-7815(B and D) consistently show a similar phenotype of outgrowth and duplication of wing pouch.

**Figure 8.**
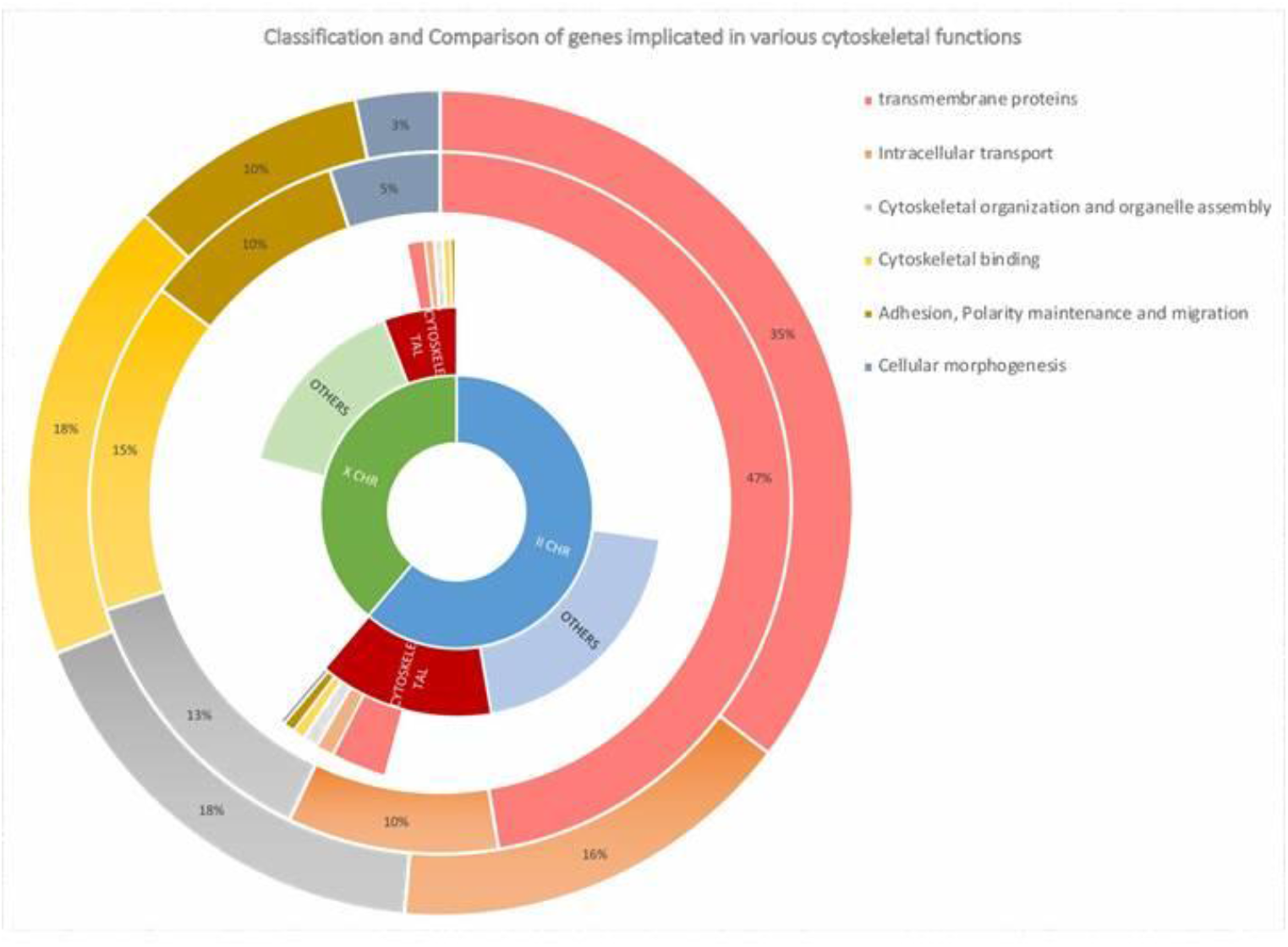
Classification and comparison of genes implicated in cytoskeletal functions. In the X chromosome, 181 genes out of 1103 total genes and in II chromosome, 431 genes out of 1245 genes are involved in mediating cytoskeletal functions. Those functions are classified into five categories and percentage of genes involved are compared and represented.

Nearly 11 genes have both fly and their corresponding human orthologs with similar cytoskeletal functions. For example, Drosophila *kirre* of Df(1)ED6330 (Fig 1F) and its human ortholog *KIRREL3* is involved in cell adhesion and plasma membrane fusion activities and the fly gene *SKPA* of Df(1)ED6443 and its human ortholog *SKP1* is implicated in microtubule cytoskeletal organization. Similarly, gene linked to Notch regulation, Drosophila *sno* of Df(1)ED7170 (Fig 1L) and its human ortholog *SBNO1* and gene connected to synaptic growth, fly *Efhc1*.*1* of Df(1)ED7344 (Fig 1I) and its human ortholog *EFHC2* are implicated in cytoskeletal morphogenesis. Few of the fly orthologs are uncharacterized while cytoskeletal functions of their human orthologs are known. For example, fly orthologs *CG10804* and *CG16181* of human genes *TROVE2* and *MARCH5* respectively are uncharacterized genes. These two genes are covered by the deficiency line Df(1)Exel6233. In the same section above we have shown that Df(1)Exel6233 produced mild cytoskeletal defects (Fig 4). Therefore it is possible that the genes *CG10804* and *CG16181* may be implicated in cytoskeletal functions. Clonal properties of 18 genes on respective deficiency lines are undertaken in this study (Supplementary data S2(Sheet1-2); Table 2). The cytoskeletal phenotypes from the fly can thus be utilized to predict the etymology of many rare, unexplored human diseases.

### 3. A Systematic approach to compare genes on Drosophila deficiency lines with Human Mendelian Diseases (MenDs) and Microdeletion Syndrome (MDs) to draw synteny between them

We undertook a systematic approach for both X and II chromosomes to compare the Drosophila genes on deficiency lines with their corresponding Human Orthologs. We have classified them into two categories based on whether or not they have available Human orthologs. There are 600 genes on X chromosome and 1090 genes on II chromosomes with Human orthologs. These groups with Human orthologs were looked up for their association with disease enlisted in Online Mendelian Inheritance in Man (OMIM) database. There are nearly 189 genes on X chromosome and 509 genes on II chromosome with OMIM disease association. Based on this criterion, we further categorized the diseases into Mendelian disease and Microdeletion Syndrome. There are nearly 119 genes on X chromosome and 227 on II chromosome for Microdeletion syndrome (Figure 9(A-B), Supplementary data S3 (Sheet1-2)) and 70 genes on X chromosome and 202 genes on II chromosome associated with Mendelian Disease (Figure 9(A-B), Supplementary data S3(Sheet 7)).

**Figure 9.**
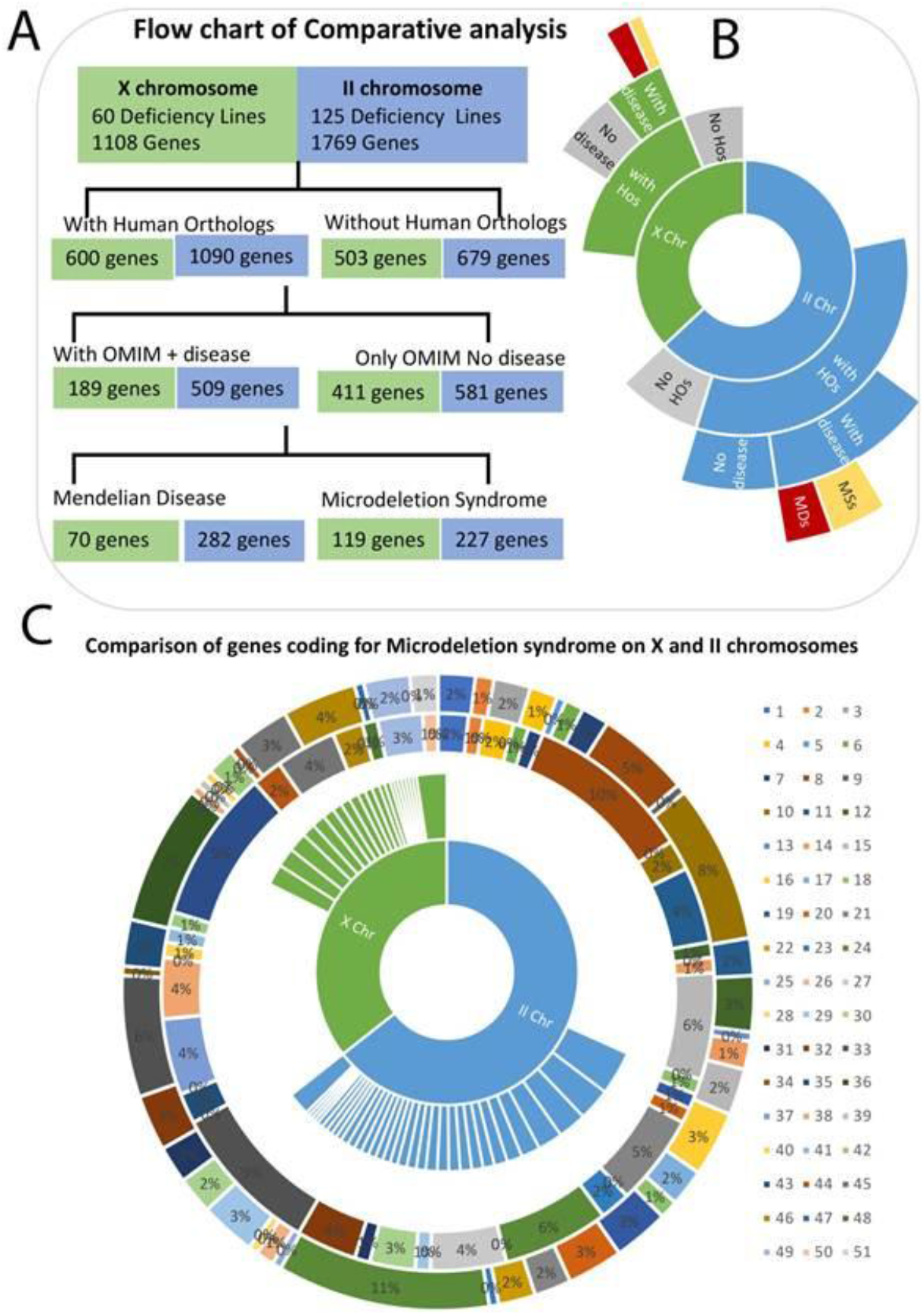
Comparative analysis of Drosophila deficiency lines with Human Mendelian Diseases and Microdeletion Syndrome. A detailed Flowchart (A) and a sunburst representation(B) of systematic approach to compare genes in the Drosophila Deficiency lines of X and II chromosome with human orthologs . Analytical representation of comparison of genes on X and II chromosomes coding for 52 human Microdeletion syndrome (c).

Further we have classified and analysed the data in two ways. First, for any locus on the human chromosome, we have identified all possible MDs or MenDs pertaining to it. Therefore, any lesion on that particular locus can lead to any of these disorders listed Supplementary data S3 (Sheet1-2), (Sheet 7)). Some of the MDs are coded by more than one locus on either of these chromosomes while some of them are either present on X or II chromosome. We have pooled and compared the number of genes on both X and II chromosome that represent the corresponding MD syndrome to generate a comprehensive comparative chart between them (Table 3, Figure9C).

**Table 3:** Deficiency lines in Drosophila and Microdeletion Syndromes in Humans

| **Microdeletion_Syndromes** | **<u>X</u>** | **<u>Ii</u>** | **Prevalence in Humans** | **Mutation hotspot in Humans** |
| --- | --- | --- | --- | --- |
| 17q21.31 microdeletion syndrome | 2 | 4 |  | KAT8 regulatory NSL complex subunit 1 - KANSL1 |
| 1p36 deletion syndrome | 1 | 2 |  | NA |
| 1q21.1 deletion syndrome, 1.35 Mb | 0 | 4 |  | GJA5, GJA8 (CHD1L, BCL9, and PRKAB2) |
| 3q29 microdeletion syndrome | 2 | 3 |  | NA |
| Adrenal hypoplasia, Congenital | 0 | 1 | 25% | NROB1, GK, DMD, IL1RAPL1 |
| Angelman's syndrome | 1 | 2 |  | MBD5 |
| Aniridia | 1 | 3 | 17-22% | PAX6 |
| Autism | 11 | 10 |  | ADNP |
| Basal Cell Nevus syndrome | 0 | 1 | 1/6-21% | PTCH1, SUFU |
| Beckwith-Wiedemann Syndrome | 2 | 17 |  | H19, IGF2, CDKN1C, KCNQ1, KCNQ1OT1 |
| Brachydactyly mental retardation syndrome | 5 | 4 |  | HDAC4 |
| Branchiootorenal syndrome | 1 | 6 |  | EYA1, SIX5, SIX1 |
| Bruton agammaglobulinemia | 0 | 1 | 8% | BTK |
| Campomelic Dysplasia | 1 | 3 | rare | SOX9 |
| Charcot-Marie-Tooth Syndrome | 7 | 5 |  | GJB1 |
| Cleidocranial dysplasia | 0 | 7 | 10% | RUNX2 |
| Cornelia De Lange syndrome | 0 | 4 |  | HDAC8, NIPBL, RAD21, SMC1A, SMC3 |
| Cri-du-chat syndrome | 1 | 2 |  | catenin delta 2 - CTNND2<br>semaphorin 5A - SEMA5A |
| Dandy Walker syndrome | 1 | 6 |  | ZIC1, ZIC4 (PROBABLY) |
| Diaphragmatic Hernia, Congenital | 1 | 6 | rare | *NR2F, CHD, RGMA, DISP1, GATA4 and SIAT8B* |
| Digeorge syndrome | 6 | 4 |  | TBX1 |
| Down syndrome | 0 | 4 | rare | NA |
| Greig Cephalopolysyndactyly syndrome | 2 | 1 | 5-10% | GLI3 |
| Holoprosencephaly | 7 | 23 |  | SHH, ZIC2, SIX3, TGIF1, GLI2, PTCH1 |
| Kabuki syndrome | 0 | 1 |  | KDM6A, KMT2D |
| Leri-Weill dyschondrosteosis | 0 | 2 | rare | SHOX |
| Mental retardation Autosomal Dominant 51 | 5 | 0 |  | NA |
| Metachromatic leukodystrophy | 0 | 1 | <1% | ARSA |
| Microphthalmia, Syndromic 7 | 1 | 6 |  | COX7B, HCCS, NDUFB11 |
| Miller-Dieker Lissencephaly syndrome | 3 | 4 |  | PAFAH1B1 |
| Muscular dystrophy, Becker Type | 1 | 4 |  | DMD |
| Nail-patella syndrome | 4 | 6 | 5-10% | LMX1B |
| Neurofibromatosis | 10 | 13 |  | NF1 |
| Neuropathy hereditary, with liability to pressure palsies | 0 | 1 |  | PMP22 |
| Noonan Syndrome 1 | 2 | 5 |  | MAP2K2, NF1 (PROBABLY) |
| Polycystic kidney disease, infantile severe, with Tuberus Sclerosis | 0 | 15 |  | PKD1, TSC2 (PROBABLY) |
| Potocki-Lupski syndrome | 5 | 0 |  | FLCN, RAI1 |
| Potocki-Shaffer syndrome | 4 | 1 |  | ALX4, EXT2, PHF21A (PROBABLY) |
| Prader-Willi Syndrome | 0 | 1 |  | SIM1 |
| Retinoblastoma Rb1 | 1 | 1 | 8-16% | RB1 |
| Rett's syndrome | 1 | 0 | 8% | MECP2 |
| Rieger Syndrome Type I | 1 | 3 |  | PITX2, FOXC1 |
| Rubinstein Taybi Syndrome | 10 | 0 |  | CREBBP, EP300 |
| Saethre Chotzen syndrome | 2 | 1 |  | TWIST1 |
| Split hand/foot malformation 1 | 4 | 6 | rare | TP63 |
| Towne's Brocks syndrome | 2 | 8 | 5% | SALL1 |
| Trichorhinophalangeal syndrome | 0 | 1 | 3-4% | EXT1, RAD21, TRPS1 |
| Tuberous Sclerosis | 1 | 0 | <2% | TSC1, TSC2 |
| William-Beuren region duplication syndrome | 3 | 5 |  | ELN, NCF1 |
| Wilm's Tumor 1, Mental retardation | 1 | 0 | 14% | PAX6 |
| Wolf-Hirschhorn Syndrome | 0 | 3 |  | CPLX1, CTBP1, FGFR1, LETM1, NELFA, NSD2, PIGG |

Second, we found many human Microdeletion and Mendelian syndromes to mirror the genetic deletions observed in the fly deficiency lines under study. We could thus make comparison between them to identify etymologies of rare variants of human genetic disorders. We have analysed the MDs by comparing the order of genes as they are arranged on the human chromosomes (synteny). In this study, we have found that there are number of genes in the order on Drosophila chromosome to be represented in the same order on Human chromosome leading to either Microdeletion syndromes (Supplementary data S3(Sheet 5-6)) or Mendelian disease (Supplementary data S3(Sheet 7)). Interestingly there are also 16 Mendelian Disease genes on X chromosome of Drosophila that correspond to genes on Human X chromosome (Table 4 or Supplementary Data S3(Sheet 8)).

**Table 4:**
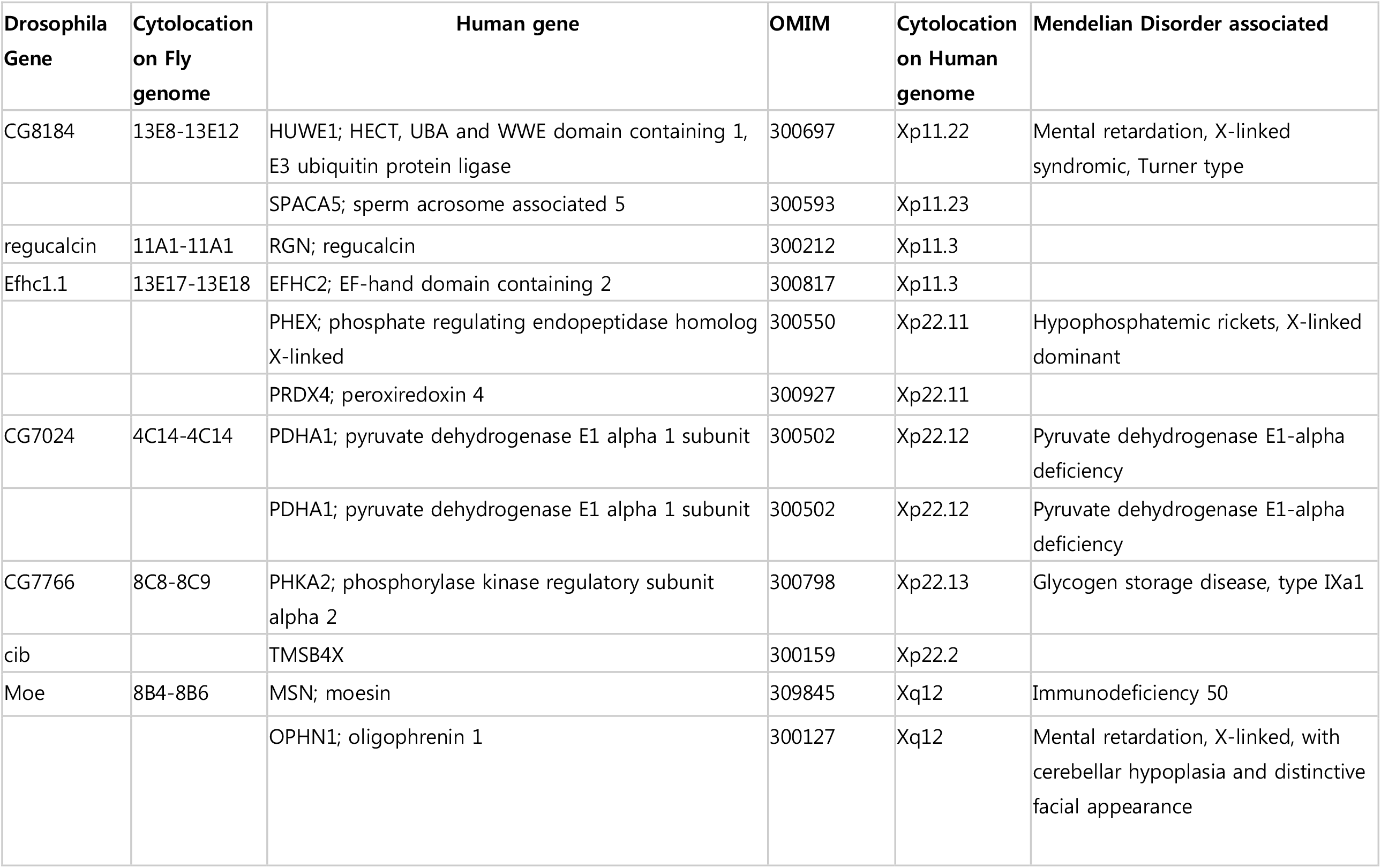
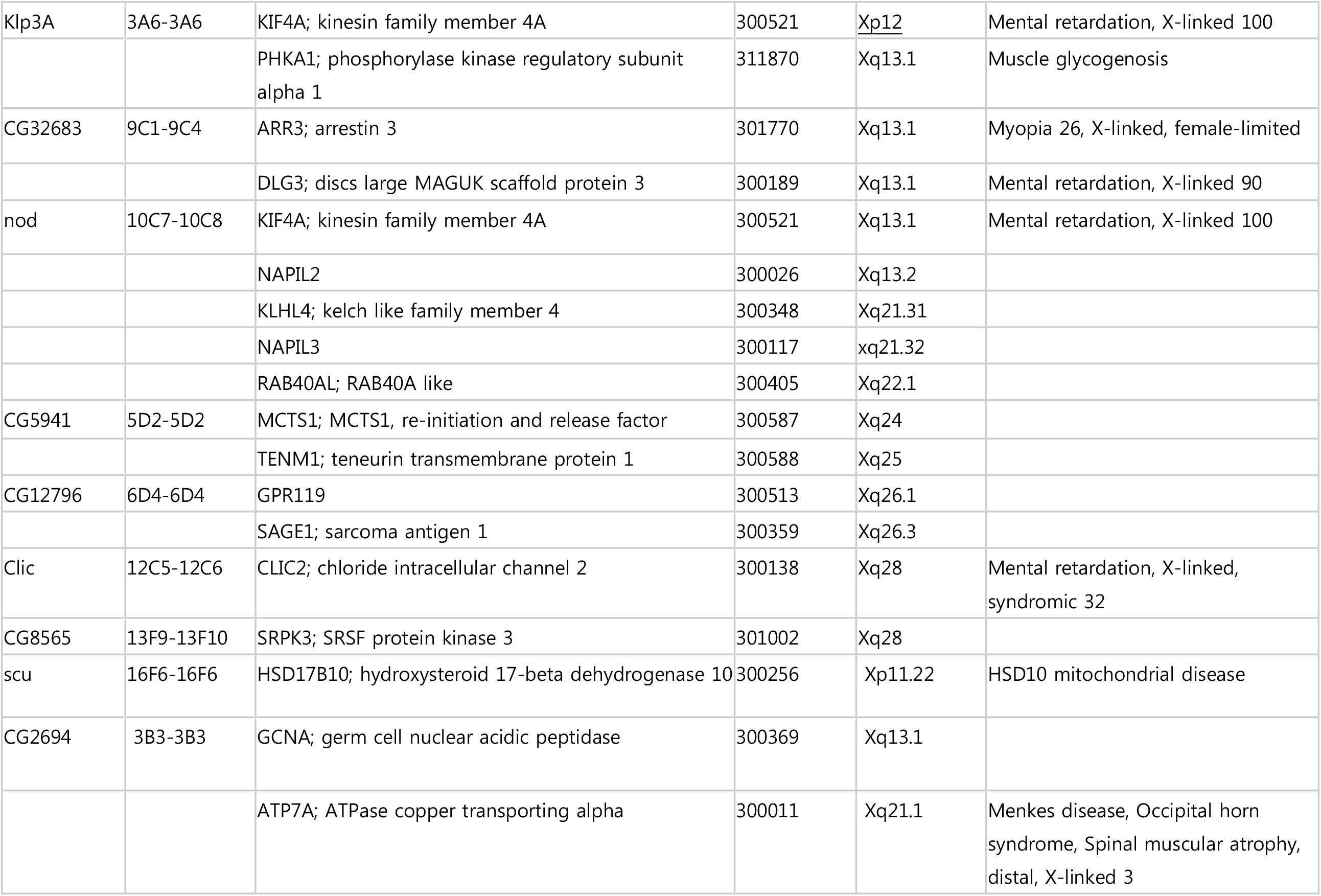
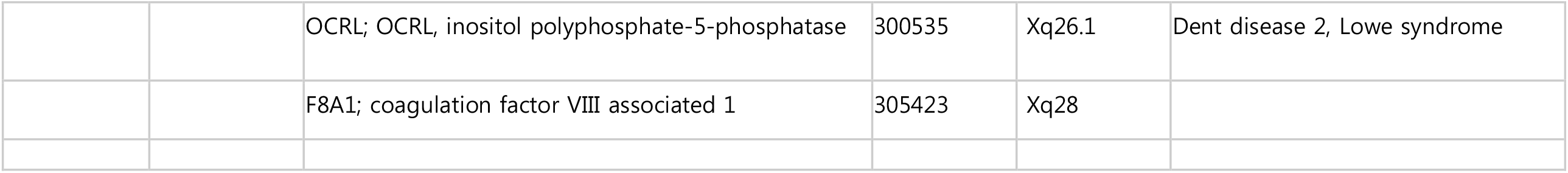
Correlation of genes on X chromosomes of Drosophila and Human associated with Mendelian diseases

| **Drosophila Gene** | **Cytolocation on Fly genome** | **Human gene** | **OMIM** | **Cytolocation on Human genome** | **Mendelian Disorder associated** |
| --- | --- | --- | --- | --- | --- |
| CG8184 | 13E8-13E12 | HUWE1; HECT, UBA and WWE domain containing 1, E3 ubiquitin protein ligase | 300697 | Xp11.22 | Mental retardation, X-linked syndromic, Turner type |
|  |  | SPACA5; sperm acrosome associated 5 | 300593 | Xp11.23 |  |
| regucalcin | 11A1-11A1 | RGN; regucalcin | 300212 | Xp11.3 |  |
| Efhc1.1 | 13E17-13E18 | EFHC2; EF-hand domain containing 2 | 300817 | Xp11.3 |  |
|  |  | PHEX; phosphate regulating endopeptidase homolog X-linked | 300550 | Xp22.11 | Hypophosphatemic rickets, X-linked dominant |
|  |  | PRDX4; peroxiredoxin 4 | 300927 | Xp22.11 |  |
| CG7024 | 4C14-4C14 | PDHA1; pyruvate dehydrogenase E1 alpha 1 subunit | 300502 | Xp22.12 | Pyruvate dehydrogenase E1-alpha deficiency |
|  |  | PDHA1; pyruvate dehydrogenase E1 alpha 1 subunit | 300502 | Xp22.12 | Pyruvate dehydrogenase E1-alpha deficiency |
| CG7766 | 8C8-8C9 | PHKA2; phosphorylase kinase regulatory subunit alpha 2 | 300798 | Xp22.13 | Glycogen storage disease, type IXa1 |
| cib |  | TMSB4X | 300159 | Xp22.2 |  |
| Moe | 8B4-8B6 | MSN; moesin | 309845 | Xq12 | Immunodeficiency 50 |
|  |  | OPHN1; oligophrenin 1 | 300127 | Xq12 | Mental retardation, X-linked, with cerebellar hypoplasia and distinctive facial appearance |
| Klp3A | 3A6-3A6 | KIF4A; kinesin family member 4A | 300521 | Xp12 | Mental retardation, X-linked 100 |
|  |  | PHKA1; phosphorylase kinase regulatory subunit alpha 1 | 311870 | Xq13.1 | Muscle glycogenosis |
| CG32683 | 9C1-9C4 | ARR3; arrestin 3 | 301770 | Xq13.1 | Myopia 26, X-linked, female-limited |
|  |  | DLG3; discs large MAGUK scaffold protein 3 | 300189 | Xq13.1 | Mental retardation, X-linked 90 |
| nod | 10C7-10C8 | KIF4A; kinesin family member 4A | 300521 | Xq13.1 | Mental retardation, X-linked 100 |
|  |  | NAPIL2 | 300026 | Xq13.2 |  |
|  |  | KLHL4; kelch like family member 4 | 300348 | Xq21.31 |  |
|  |  | NAPIL3 | 300117 | xq21.32 |  |
|  |  | RAB40AL; RAB40A like | 300405 | Xq22.1 |  |
| CG5941 | 5D2-5D2 | MCTS1; MCTS1, re-initiation and release factor | 300587 | Xq24 |  |
|  |  | TENM1; teneurin transmembrane protein 1 | 300588 | Xq25 |  |
| CG12796 | 6D4-6D4 | GPR119 | 300513 | Xq26.1 |  |
|  |  | SAGE1; sarcoma antigen 1 | 300359 | Xq26.3 |  |
| Clic | 12C5-12C6 | CLIC2; chloride intracellular channel 2 | 300138 | Xq28 | Mental retardation, X-linked, syndromic 32 |
| CG8565 | 13F9-13F10 | SRPK3; SRSF protein kinase 3 | 301002 | Xq28 |  |
| scu | 16F6-16F6 | HSD17B10; hydroxysteroid 17-beta dehydrogenase 10 | 300256 | Xp11.22 | HSD10 mitochondrial disease |
| CG2694 | 3B3-3B3 | GCNA; germ cell nuclear acidic peptidase | 300369 | Xq13.1 |  |
|  |  | ATP7A; ATPase copper transporting alpha | 300011 | Xq21.1 | Menkes disease, Occipital horn syndrome, Spinal muscular atrophy, distal, X-linked 3 |
|  |  | OCRL; OCRL, inositol polyphosphate-5-phosphatase | 300535 | Xq26.1 | Dent disease 2, Lowe syndrome |
|  |  | F8A1; coagulation factor VIII associated 1 | 305423 | Xq28 |  |

Until now, researchers and clinicians were finding it difficult to pin down the respective genes that correspond to the disease manifestation. However, from this comprehensive analysis, it is possible to identify the specific gene that lead to MDs and MenDs. For example, we found that one of the MDs, Holoproscencephaly, is caused by loss of several genes. Nearly there are 7 genes on X and 23 genes on II chromosome that can lead to the manifestation of the syndrome (Table 3, Supplementary Data S3 (Sheet1-2). Next highest are the MDs like Autism (11 genes on X and 10 genes on II chromosome) and Neurofibromatosis (10 on X chromosome and 13 on II chromosome). Comparative loci analysis reveals that in the locus on human 2q37.3, either Autism or Holoprosencephaly can occur. By comparing with the Drosophila genes, it is possible to point out the genes and locus on the chromosome. While loss of *per* (at 3B1-3B2) can lead to Autism, loss of *CG10353* (at 10F2-10F4) can lead to Holoproscencephaly Supplementary Data S3 (Sheet1, Sheet3). Similarly, lesions at loci 11p11.2 or 16p11.2 can lead to DiGeorge Syndrome/Velocardiofacial syndrome and Holoproscencephaly. From this study we find that loss of either *SPACA3* at 11p11.2 or *FAM57B* at 16p11.2 can lead to DiGeorge Syndrome, while loss of either *PTPRI* at 11p11.2 or loss of *PHKG2* at 16p11.2 can lead to Holoproscencephaly (Supplementary data S3(Sheet1, Sheet3)).

Similarly, the locus 5A12-5D1 encodes for several essential genes and therefore any deficiency in this region may potentially give rise to MDs like Beckwith Wiedemann Syndrome (BWS), Bracydactyly-Mental Retardation syndrome (Bdmr1), (Branchiorenal syndrome, Bor1 and Campomelic Dysplasia). Deficiency line Df(1)ED6802(BL-8949), pertaining to the locus 5C2-5C2, that encodes for Carboxy peptidases A3 and B1 (CPA3) may lead to BWS as well as for Bdmr1. This also suggests that Human orthologs of these genes could potentially be involved in human X-linked MDs) (Supplementary data S3(Sheet1, Sheet3)).

Interestingly, we have found that there are MDs that are caused by loss of genes only on X chromosome and not on II chromosome. For example, Rubinstein Taybi Syndrome is caused by 10 loci, all located only on X chromosome and not on II chromosome. Similarly, Potocki-Lupski Syndrome is caused by 5 loci on X chromosome and not on II chromosome. Rett’s syndrome and Tuberus sclerosis are MDs caused by microdeletions in *MECP2* gene and *TSC1/2* respectively both with single locus each on X chromosome. (Table3, Supplementary data S3(Sheet1, Sheet3)). Corresponding deficiency lines analysed here are Df(1)ED6630-8948 (Figure 1E) and Df(1)ED7170-9058(Figure 1K) that has the genes *CG2680* and *HDAC4* respectively representing the microdeletions causing Rett’s syndrome and Tuberus sclerosis.

From this analysis, in addition to the point that a particular MDS can be represented by more than one locus, we also understand that, multiple MDs that arise in a particular locus depend on the genes deleted and the order of contiguous genes deleted. In addition, comprehensive and comparative analysis between Human and Drosophila loci have led to identification of putative critical region in human chromosome that could give rise to various MDs. This is addressed in detail in the following section.

### 4. Identification of critical region or the gene implicated in epithelial defect

Although multiple genes are lost in the manifestation of MDs or Mendelian diseases, there exists a critical region involving either an essential gene or a small genomic region in that locus which is central in causing the disease. Since the molecular ends of the deficiency lines used in the study were precisely defined beforehand, we compared this information with the clonal phenotype or behaviour generated to identify the critical region in that locus, necessary for the maintenance of tissue integrity. We took two approaches while comparing the overlapping deficiency lines, one based on the number and size of the clones generated with respect to the number of genes eliminated (Supplementary S1 Sheet(2 and 4); and the other based on the cytoskeletal defects displayed by them (Supplementary S3). For the former approach, using the deficiency lines on X chromosome, we looked into the category with numerous clones (>60 clones per wingdisc) generated by a smaller number of genes lost in them. Of the 10 deficiency lines in this category, deficiency line Df(1)Exel6237 (BDSC-7711)at 5C2-5C6 with 21 genes lost including *Usp30* generated 103 clones, the highest number of clones of all lines. Followed by that, Df(1)Exel6230(BDSC-7705) at 3A2-3A4 with 9 genes lost generated biggest clones(with average size of 0.6 Sq.pixels) and the second highest number of clones per wing disc (on an average 91 clones). Under this category, we also found an overlapping deficiency line Df(1)Exel6231(BDSC-7706) for the same region at 3A2-3A3, that lost 6 genes but produced 65 clones (6^th^ highest number of clones) with average size of 0.4 Sq.pixels(one of the biggest clones). For the same locus, we also found two bigger deficiency lines Df(1)ED11354 (BDSC-9345) at 2F6-3A4, with 15 genes lost, but generating only 13 clones (with average size of 0.05 Sq.pixels); and Df(1)ED411(BDSC-8031) at 3A3-3A8 with 18 genes lost, but produced only 3 clones on an average(with average of size 0.2 Sq.pixels). Of the four deficiency lines spanning this locus, (Df(1)Exel6230(BDSC-7705) and Df(1)Exel6231(BDSC-7706) show similar phenotype with large (0.6 and 0.4 Sq.pixels)and numerous clones (91 and 65) respectively. By comparing these three deficiency lines (Df(1)Exel6230(BDSC-7705), Df(1)Exel6231(BDSC-7706) with Df(1)ED411(BDSC-8031), we found that 3A2-3A3, that contains the gene *Brother of ihog (Boi)* is the critical region/gene responsible for producing high number of bigger clones (Figure10A, 10E and 10H ; Supplementary Data S4 (Sheet1). It is understood that the third biggest deficiency line Df(1)ED11354 (BDSC-9345) might include other genes like *Raf*, another regulator of growth. *Boi* is suggested to be a negative regulator of growth, that functions as a co-receptor for Patched(Ptc) in regulating Hedgehog(Hh)-signaling mediated growth. Therefore, highest number of clones generated is due to the elimination of *Boi*, the negative regulator of growth, in that locus. Interestingly, loss of the gene *Cell adhesion associated, Oncogene regulated (CDON)*, human ortholog of Boi, localized at 11q24.7 is suggested to cause MD Holoprosencephaly in humans.

As a second approach, on both X and II chromosomes, we looked into those deficiency lines which generated cytoskeletal defects and wingdisc phenotypes associated with them. On X chromosome, the deficiency line Df(1)ED6584(BDSC-9348) (Figure11C) at 3A8-3B1 lost only 3 genes, but generated moderate number of clone (average number of 36 and average size of 0.12 Sq.pixels). We compared it with available overlapping deficiency lines Df(1)ED6679 (BDSC-9518) (Figure11(A’-A”)) at 3A6-3A8 with 10 genes lost, that produced only 8 clones (with average size of 0.08 Sq.pixels); and Df(1)ED411(BDSC-8031) (Figure11B) at 3A3-3A8 with 18 genes lost produced only 3 clones on an average(with average of size 0.2 Sq.pixels). The clonal analysis of the deficiency line Df(1)ED6679 (BDSC-9518) by both conventional method Figure 11 A-A’) and MARCM technique (Figure11(D’-D”)) showed several cytoskeletal defects in the hinge region, with small rounded cells around the periphery of the hinge. However, the clonal analysis of Df(1)ED6584(BDSC-9348) showed cell-autonomous extrusion of the clones from the wing surface (Figure11(E’-E”)). By comparing the overlapping region between them and the similar phenotype produced by Df(1)ED6679 (BDSC-9518), we could identify the critical region is coded by the gene *Shaggy(Sgg)*. As *Sgg* is a key component of β-catenin destruction complex so as to function as a negative regulator of Wg-signaling pathway and Insuline-like receptor signaling pathways, we corroborate that the phenotype and the critical region involved in these deficiency lines is encoded by *Sgg* (Figure10B, 10F, Figure11C, Figure11(D‘-D”).

**Figure 10.**
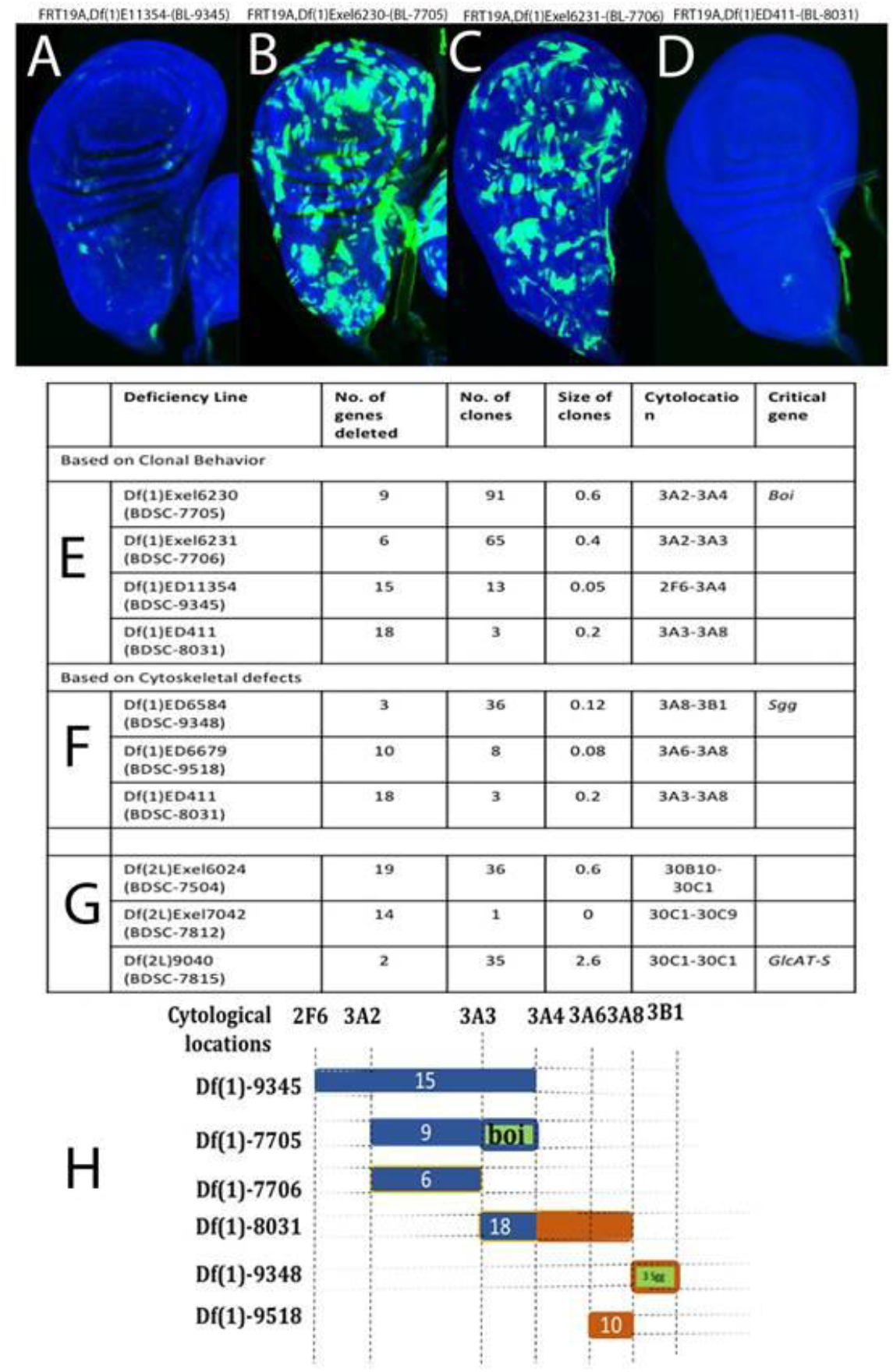
Identification of critical region based on clonal behaviour. For the locus 2F6-3A8, two big deficiency lines BL-9345 (10A, E) and BL-8031(10D-E) produced a smaller number of clones while two smaller deficiency lines BL-7705 (B, E) and BL-7706(C, E) produced high number of big sized clones (B, E). By comparing their clonal properties and cytoskeletal information, we identify that the critical gene is *Boi* (E). Similarly, for the loci 3A3-3B1 and 30B10-30C9 (for clones Fig 11, Fig12) the critical gene is *Shaggy (Sgg)* and *GlcAT-S(F)* respectively. Schematic representation of the critical region identified in this study (H).

**Figure 11.**
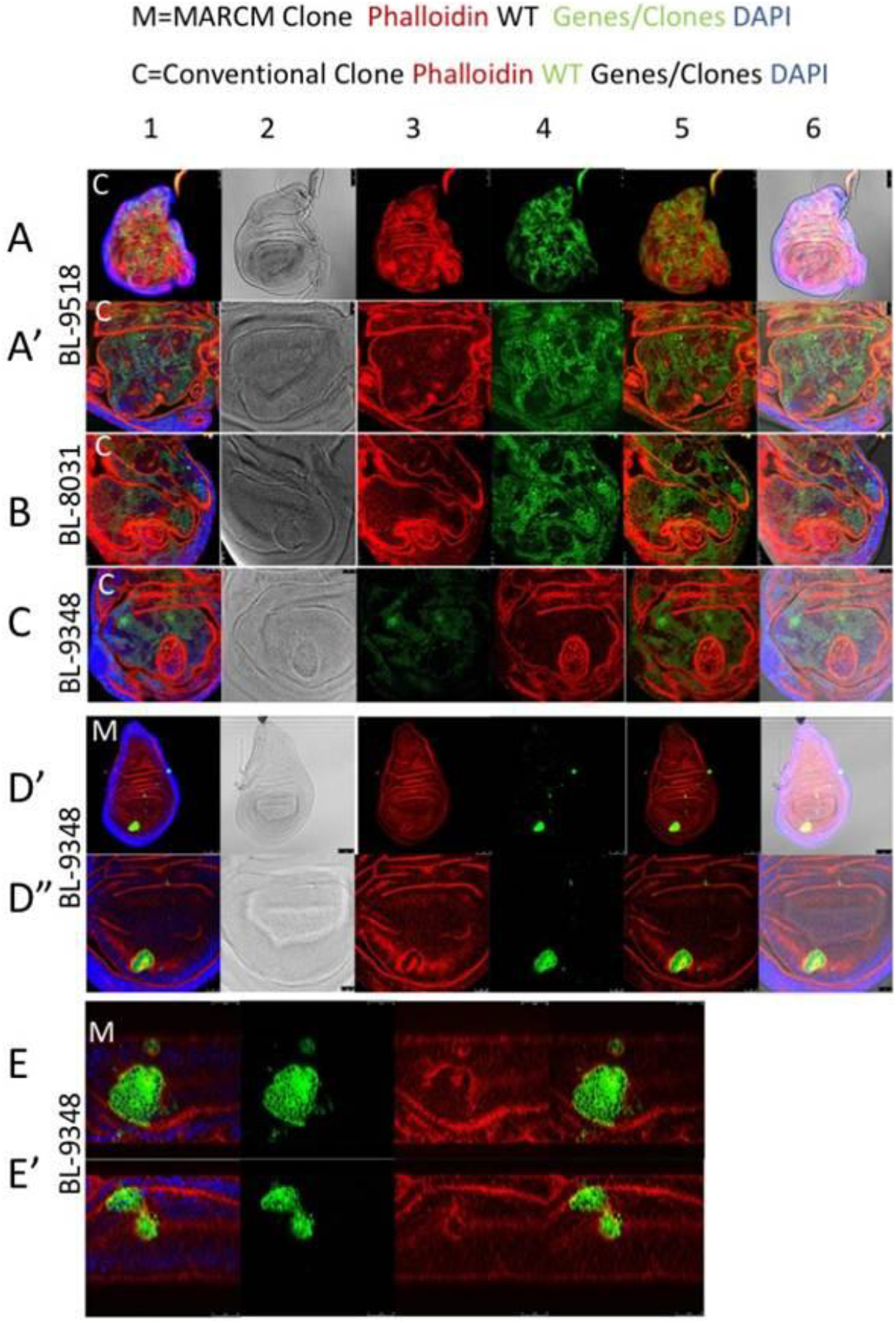
Identification of critical region based on the cytoskeletal defects. On the X chromosome, three deficiency lines BL-9518(A-A’), BL-8031(B) and BL-9348(C) that produced moderate number of clones were compared for their cytoskeletal defects. For the locus 3A3-3B1, severe phenotype produced by BL-9348(C, D’-D”) showing cell-autonomous extrusion of the clones (C, D’-D”, E-E’) from the wing surface is encoded by the critical gene *Shaggy (Sgg)*.

Similarly we have compared all other overlapping deficiency lines that produced cytoskeletal defects and identified the possible critical regions responsible for the phenotype (Figure10). The deficiency line Df(1)Exel6242(BDSC-7716) localized at 10D1-10D7 showed outgrowth and protruding phenotype of the wing disc (Figure 12B). When we looked for overlapping regions, we found the deficiency line Df(1)ED7067 (BDSC-9154) localized adjacent to it but not completely overlapping. Therefore we generated an FRT-recombination for a smaller region using a deficiency line Df(1)Exel9050(BDSC-7759), losing 3genes at 10D5-10D6. The cytoskeletal defect produced was similar to that produced by the deficiency line Df(1)Exel6242(BDSC-7716) (Figure 12(B-C)) and therefore we can suggest that the critical region for the epithelial defect is because of the gene *α1,6-fucosyltransferase (FucT6)* localized to 10D5-10D5. *FucT6* is an uncharacterized gene suggested to be involved in N-glycan Fucosylation and N-linked glycosylation (Roos et al., 2002; Paschinger et al., 2005; Frappaolo et al., 2018). Similarly for overlapping regions, using deficiency lines Df(1)Exel6226(BDSC-7703) at 1E3-1F3 that generated on average only 12 clones and Df(1)ED6521(BDSC-9281) at 1E3-1F4 that generated 46 clones on an average, we have identified the critical region encoding a previously uncharacterized gene as *N(alpha)acetyltransferase 30A (Naa30A)* localized at 1F3-1F4. Along the same lines, for deficiency lines Df(1)Exel6234(BDSC-7708) at 4F10-5A2 that generated on an average 42 clones and Df(1)Exel6235 (BDSC-7709) at 5A2-5A6 that generated 64 clones, we have narrowed it down on an uncharacterized gene *CG12730* localized at 5A2 to the critical gene/locus. There are few more overlapping deficiency lines, but since the critical gene identified further included more than two or three genes, and due to unavailability of stocks to generate clones, we were unable to characterize them in this present study. Using the same approach, for II chromosome, we compared two overlapping deficiency lines that produced opposing phenotypes to each other. The Deficiency line Df(2L)Exel7042(BDSC-7812)(Figure13B) at 30B10-30C1 that lost 14 genes produced 36 clones while an overlapping deficiency line Df(2L)Exel6024(BDSC-7507) (Figure13C) at 30C1-30C9 that lost 19 genes produced clones very rarely(No clones to 1 clone). Interestingly when we compared the wingdiscs in both cases, we found them to be slightly bigger than the control wings (Figure13(A2,B2,C2)). Therefore we took another overlapping deficiency line Df(2L)9040(BDSC-7815) available around 30C1-30C1. On generating clones, it produced overgrowth and duplication of wingdiscs(Figure13(D’-D”)). Since there were two genes, *Thioredoxin(Trx2)* and *Glcuronyl Acetyl Transferase-S (GlcAT-S)*in that region, we generated overexpression clones for both of them. We found that overexpression of *GlcAT-S* clones and not *Trx2* (data not shown), produced cytoskeletal defects ((Figure13E). We also used a P-element insertion at *GlcAT-S* that functions as a hypomorph, recombined with FRT chromosome *(FRT40A-GlcAT-SKG)* for clonal analysis. We found that GlcAT-S^KG^ reproduced the overgrowth and cytoskeletal defects(Figure13G), indicating that *GlcAT-S* localized at 30C1-30C1, is the critical region causing the epithelial defect. *GlcAT-S* is an uncharacterized gene, suggested to function as a co-receptor for glypicans and glycoproteins in regulating growth (Nagarajan et al., 2015).

**Figure 12.**
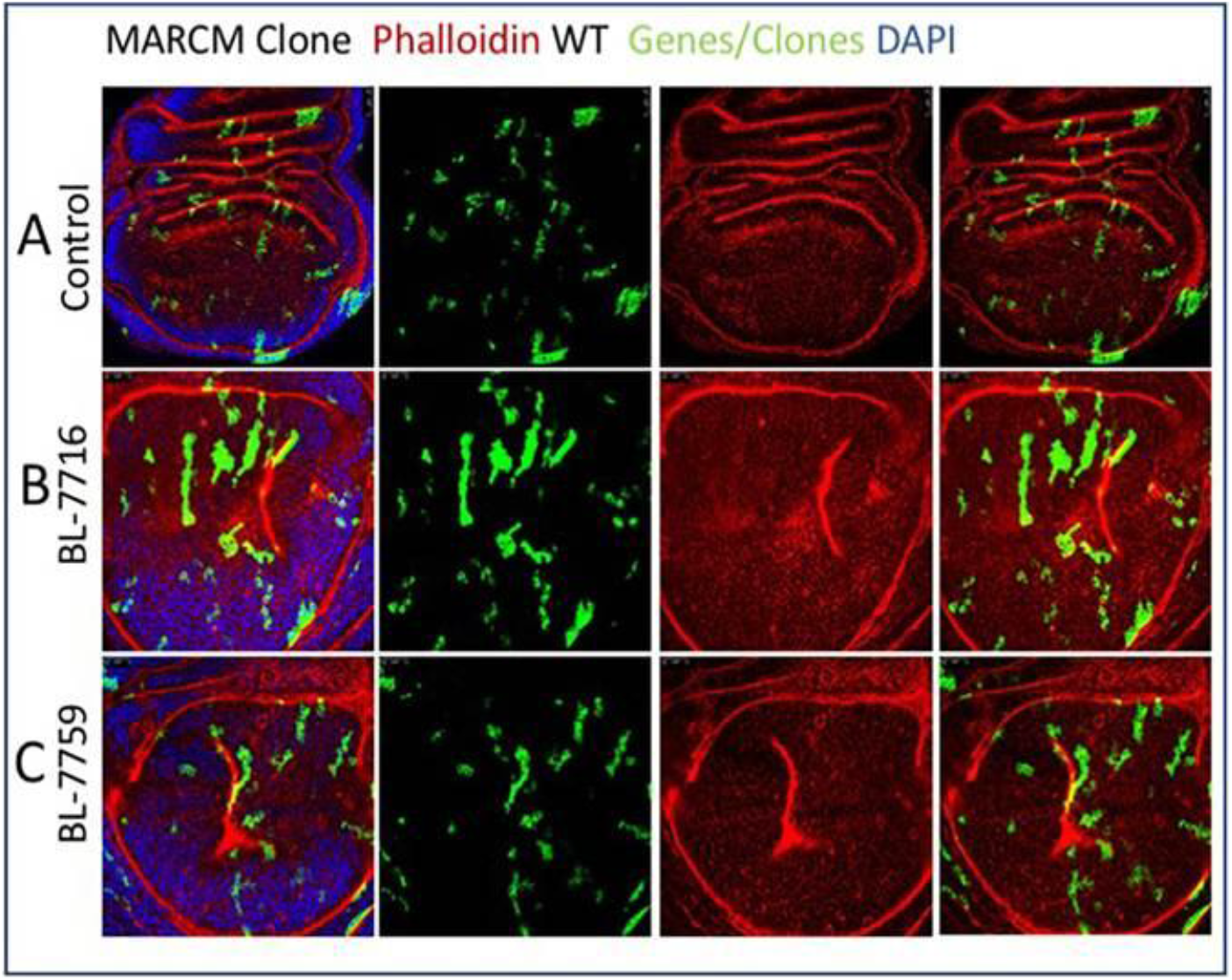
Identification of critical region using non-overlapping deficiency lines. MARCM clones for the locus 10D1-10D7, a big deficiency line BL-7716 produced a cytoskeletal defect (B). A smaller deficiency line, BL-7759 covering a gene *-1,6, fucosyltransferase (FUCT6)* produced a similar cytoskeletal defect (C).

**Figure 13.**
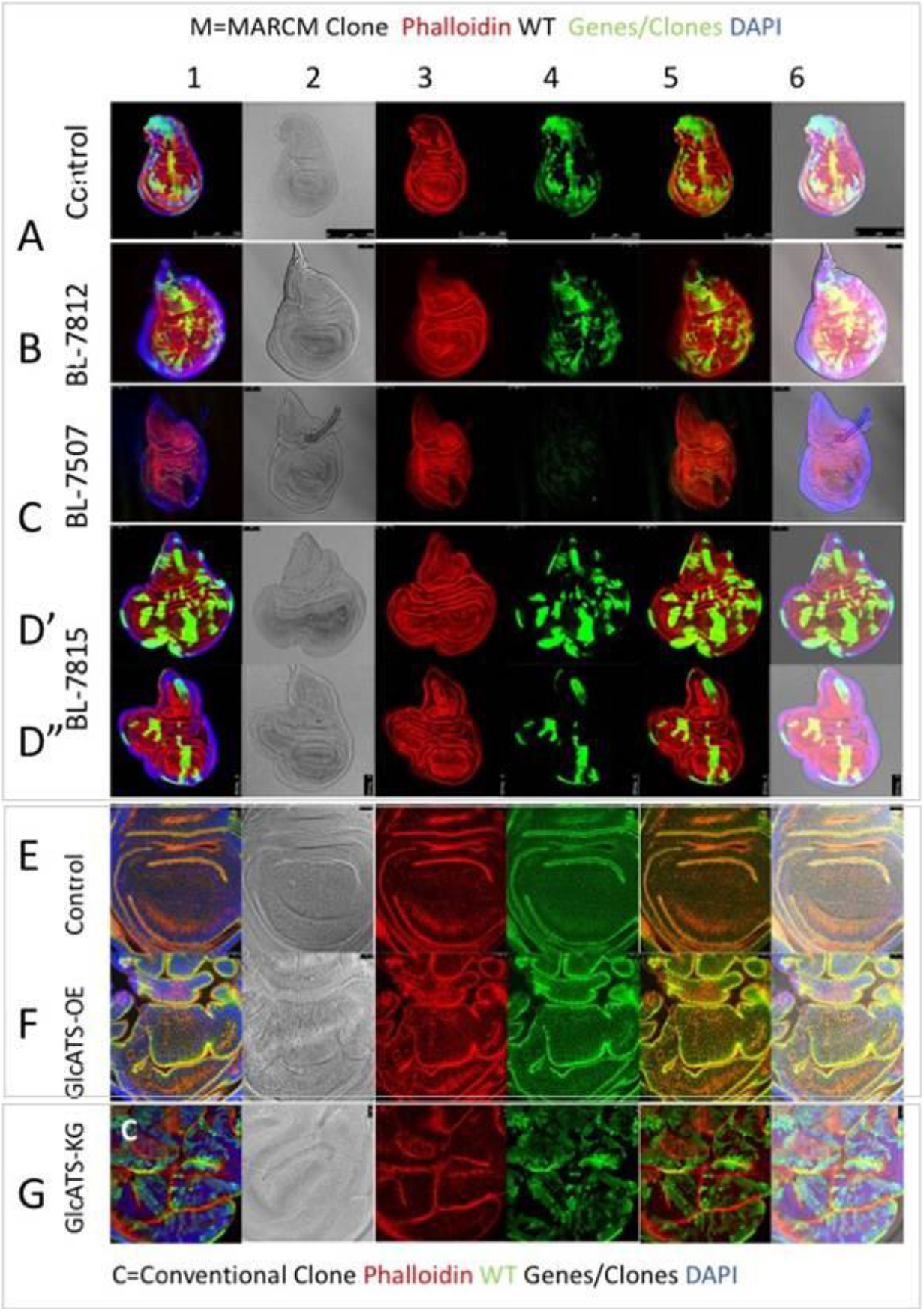
Identification of critical region for locus with opposing phenotypes For the locus 30B10-30C9, a big deficiency line (BL-7812) produced moderate number of clones (B) while an overlapping deficiency lines produced rare or very few clones (C). Note wingdiscs produced by both deficiency lines were slightly larger than the control(A-C). A small, overlapping deficiency line (BL-7815) containing two genes GlcAT-S and Trx-2 for the locus 30C1-30C1 prodeuced overgrowth and duplication of the wingdisc (D’-D”). Overexpression clones of GlcAT-S, produced cytoskeletal defects(F) and clones generated by FRT-recombined GlcAT-S^KG^ hypomorph produced overgrowth and cytoskeletal defects(G).

### Concluding Remarks

Surprisingly genetic loci of chromosomes, for instance Y chromosomes of Humans, which are restricted from recombination and spurious exchange of genetic material have been shown to undergo multigenic deletions/duplications that leads to polymorphism in the population. Though it has been intriguing to understand the consequence of such random loss of genetic material, it has remained challenging. To address this, we have utilized the fly deficiency lines. Collections of deletion-bearing Fly stocks are powerful tool utilized for mutagenesis and gene mapping. To make them even more useful, as a pilot attempt, here we have generated a novel deficiency kit comprising small-chromosomal deletion recombined onto FRT chromosomes. This collection currently comprises most of the X (∼67) and partial II chromosome (∼125) FRT-recombined deficiency lines. These stocks will be submitted and made available at the Bloomington Drosophila stock centre. In this study we have utilized these FRT recombined -deletion-bearing chromosomes for mosaic analysis in III instar wing disc to screen predominantly for cytoskeletal changes implicated in epithelial morphogenesis. In addition, researchers can also utilize this approach for any of the individual deficiency lines not included in this collection and a similar work can be extended either to other tissues for clonal analysis or to understand any other cellular processes like neurogenesis, growth processes, polarity-regulation etc. This approach helps to understand how genes behave if they are deleted in a group or when small region containing genes in order is lost. We found that overlapping deficiency lines spanning the chromosome with varied number of genes lost in them, produced clones of varied number and sizes. Clonal properties (size, number and distribution) and clonal behaviour (interaction with its neighbouring cells) of mitotic clones generated by a gene loss reflects its function. Therefore, we have further categorized them into groups based on their clonal properties to have better understanding of the gene function. Further, using these overlapping deficiency lines that generate cytoskeletal defects with phenotypes, we have identified few essential loci and genes critical for maintaining tissue integrity and growth. We also understand that those deficiency lines that do not generate clones suggest that these loci may be a crucial region encoding essential genes. As these deletion-bearing chromosomes are equivalent to human chromosomes with microdeletions, we compared the such regions that caused microdeletion-syndromes and other Mendelian disorders in humans with respective cytological locations on Drosophila chromosomes. As contiguous-gene deletion syndromes display extremely variable clinical symptoms depending on the genes lost in the loci, the existing problem in diagnosing patients with MD/MenD or other Contiguous-gene deletion syndromes or multigenic deletion syndromes is the difficulty in identifying the critical loci for disease manifestation. This study is an example to show how data from genomic information from multigenic syndrome patients or MD or MenD patients can be pooled to identify the critical loci implicated in disease manifestation. By correlating the genotype-phenotype data available for multigenic deletion syndromes and genomic information it might be possible to provide a better prognosis of the disease development.

## Experimental Procedure

### 1. Clonal Analysis

#### (i) Drosophila Stocks

Drosophila stocks and crosses were maintained on a standard yeast/cornmeal/corn syrup/malt extract/agar medium at 25°C, uncles indicated otherwise. Deletion-bearing stocks/ Deficiency lines of Exelixis, Inc (Exel), nearly ∼208 fly stocks covering X chromosome and second chromosome were obtained from Bloomington Drosophila Stock Centre. Other stocks used for this study include (5130)-pin/cyo; UAS-mCD8GFP, (5131)-hs-FLP; FRTG13, UAS-mCD8GFP, (5134)-hs-FLP, tubP-GAL80, FRT19A; UAS-mCD8GFP, (5074)-FRT40A; T155GAL4, UAS-FLP, (5075)-FRTG13; T155GAL4, UAS-FLP, (5192)-FRT40A, tubP-GAL80; (5132)-hs-FLP, tubP-GAL80, FRT19A; (5140)-FRTG13, tubP-GAL80; (5138)-tubP-GAL4; (8862)-hs-FLP; (5136)-UAS-mCD8GFP (X); (5130)-UAS-mCD8GFP (III); (42725)-hs-FLP, UAS-mCD8GFP; FRT40A, tubP-GAL80; tubP-GAL4. For generating clones by conventional method, FRT19A ; T155GAL4, UAS-FLP, FRT40A; T155GAL4, UAS-FLP and FRTG13; T155GAL4, UAS-FLP were used.

#### (ii) Recombination and Generation of clones

Sequence-Mapped deletion-bearing fly stocks (Deficiency lines) for X and II chromosomes generated by DrosDel Project and Exelixis collection (Exel) (Ryder et al., 2004; Parks et al., 2004; Cook et al., 2012; Roote and Russell, 2012) obtained from Bloomington Drosophila Stock Centre (BDSC) were recombined with their corresponding FRT chromosomes (FRT19A, FRT40A and FRTG13 chromosomes) and recombined stocks were selected on neomycin (G418) containing media. We have successfully recombined ∼67 X chromosome deficiency lines and 125 second (2L and 2R) chromosome deficiency lines with their respective FRT chromosomes. List of Deficiency lines with the number of genes lost in the chromosome is provided (Supplementary data S1-Sheet-1). Further FRT-Recombined deficiency lines were subjected to mitotic clonal analysis using both conventional and Mosaic Analysis with Repressible Cell Marker (MARCM) techniques (Lee and Luo, 1999, Lee and Luo, 2001; Blair, 2003; Wu and Luo, 2006) (Supplementary Figure1) in the III instar wing imaginal discs (Figure 1(A-O)).

Analysing clones by conventional method was tedious, therefore we followed clonal analysis by MARCM method. Total number of clones and size of each clones per wing imaginal discs were counted and recorded using ImageJ software. Nearly 10 wing discs (n=10) were imaged and replicates of three wing discs were randomly chosen for averaging total number of clones and size of each clone per wing imaginal discs.

Following protocol by Wu and Luo, 2006, MARCM Stock FRT19A (hs-FLP, tubP-GAL80, FRT19A; tubP-GAL4; UAS-mCD8GFP), MARCM Stock FRT40A (hs-FLP, UAS-mCD8GFP; FRT40A, tubP-GAL80; tubP-GAL4) and MARCM Stock FRTG13 (hs-FLP, UAS-mCD8GFP; FRTG13, tubP-GAL80; tubP-GAL4) were assembled using above stocks to generate MARCM clones. Individual FRT-recombined deletion-bearing stocks were crossed with corresponding MARCM stocks. On the 3^rd^ day AEL, first instar larvae were heatshocked at 37°C for 1hour and allowed to recover at 25°C after that. On 6^th^ day AEL, from each mutant type generated, third instar larvae were dissected and subjected to immunostaining (Usha and Shashidhara, 2010; Szuperak et al., 2011).

#### (iii) Immunostaining and Clonal Analysis

Wing Imaginal discs were immunologically stained using Phalloidin followed by Alexa594 and viewed under the confocal microscope. Their images were then analysed for clone size, distribution and number using Image J software. The clones of each of the genes mutated were analyzed in triplicates each for both the fly lines. The average clone sizes and clone numbers were calculated in each case, with the most approximate result in each case (For discs with <10 clones, all clones were considered for mean clone number estimation; For discs with >50 clones, 50 of the clones were taken into consideration for mean estimation; as well as for discs with intermediate number of clones, suitably, either 10, 20, 30 or 40 clones were considered for best estimate of the average). A triple blind test was also performed to corroborate each of these results.

### (2) Systematic Analysis

We began the systematic analysis by identifying the genes deleted in each of the available deficiency lines from http://bdsc.indiana.edu/. We then built an inventory with the deleted genes arranged according to their cytological location, ensuring that there are no repetitions, against their respective deficiency lines. Alongside, a list of their corresponding human orthologs was generated using the gene summaries available at www.flybase.org → Orthologs → Human orthologs.

Apart from this, we assessed each gene for cytoskeletal implications from the gene ontology summaries updated on their respective flybase web pages and we have categorized based on their function. (www.flybase.org → Gene Ontology → Molecular Function / Biological Processes / Cellular Component). Similarly, we also scrutinized the human orthologs for any cytoskeletal implications, under the same categories, using the database available at https://www.ncbi.nlm.nih.gov/gene/. For those genes that were involved in functions that fell into more than one of these categories, we took their most predominant function into account for grouping and analyzing.

We used the Molecular Genetics Laboratory group, MGenoma®’s webpage as a reference for a list of microdeletion syndromes. From the table present, we enlisted the genes that were deleted in each of the syndromes. We identified among them, those human orthologs that we previously enlisted in the inventory generated, and also their respective fly counterparts. For each of the microdeletion syndromes, we used the database from Orphanet to find the critical region, or the mutational hotspot. The name of the disease was given in the search field and the molecular genetics section of the clinical genetics review for each disease was referred to, in order to identify the mutational hotspot or the critical region of the gene(s) deleted.

Supplementary Data Table S1 -Excel sheet with list of deficiency lines used for recombination. X chromosome (Sheet1) and II chromosomes (Sheet 2) (Supplementary Data Table S2) – Number of clones and average size of clones Supplementary Data Figure S3)-Mitotic clones Range from no clones to big clones

## Supporting information

Supplemental DATA S1 (Sheet 1 to 5)

Supplemental DATA S2 (Sheet 1 to 5)

Supplemental DATA S3 (Sheet 1 to 5)

Supplemental DATA S4 (Sheet 1 to 5)

## Authors Contributions

Conceptualization: UN, SP; and Methodology: UN, SP, SS, RR, NR; Investigation: UN, SP, TK; Formal Analysis: UN, SP, SS, RR, NR; Writing-UN, SP, TK; Writing-Review and Editing-UN, SP, NR, TK Funding Acquisition: UN; Supervision: UN, SP.

## Acknowledgments

We gratefully acknowledge Dr Matt Gibson and Dr Marios Georgiou for their help with fly experiments in their laboratories. We thank the Bloomington stock centers and the fly community for the generosity with reagents. We thank Prof Satish Kumar and Dr Vasanthi Dasari for helpful advice and critical comments on the manuscript. We thank Science and Engineering Research Board (SERB), Department of Science and Technology (DST), Government of India, for the grants (YSS/2014/000896) and CRG/2018/003725 to UN.

---

**Supplementary Figure 1.**
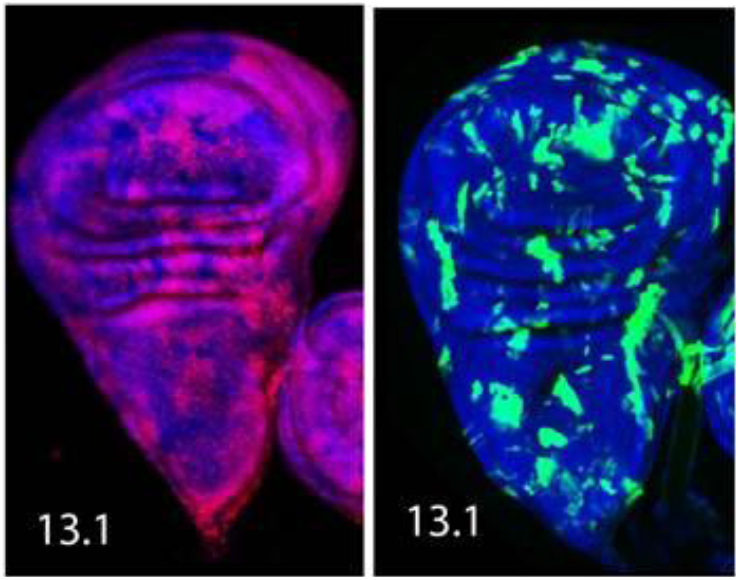
Mitotic clones. Control Wingdisc showing mitotic clones generated (A) by conventional method where clones are negatively marked, and counterstained with Phalloid (Actin, RED) and (B) by Mosaic Analysis with Repressible Cell Marker (MARCM) techniques where clones are positively marked (GFP) and counter stained with DAPI (Nucleus, Blue).

**Supplementary Figure 2.**
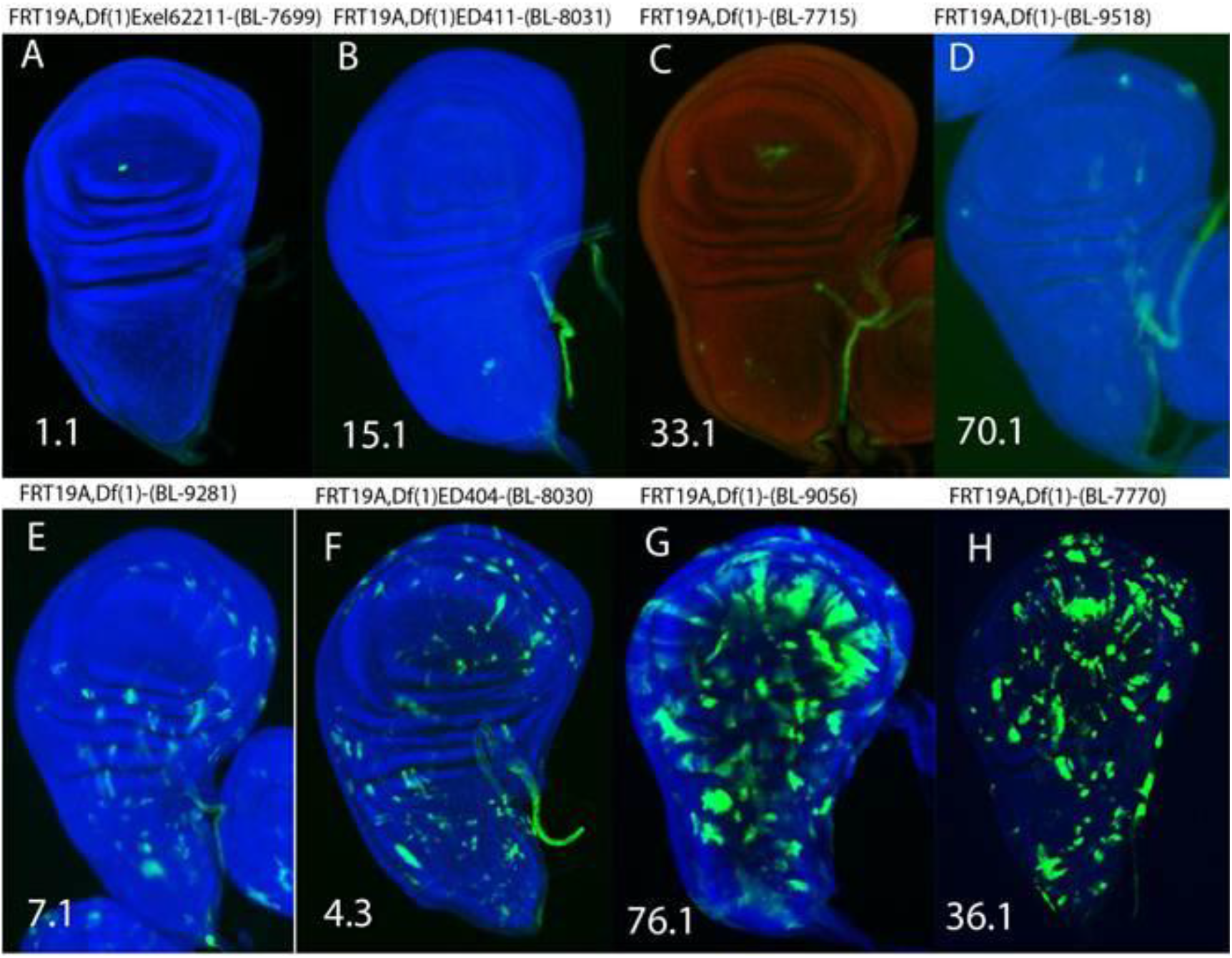
Variation in localization and number of clones. Representative Wingdiscs with localized clones (A, B, E), few clones (C, D), moderate clones (E, F) and high number of clones (G, H).

## Reference

Akiyama, T., User, SD and Gibson, MC (2018). eLife; 7: e35258. DOI: https://doi.org/10.7554/eLife.3525

Bejsovec, A. (2018). Wingless Signaling: A Genetic Journey from Morphogenesis to Metastasis. Genetics 208(4): 1311–1336.

Blair SS (2003). Genetic mosaic techniques for studying Drosophila development. Development 130: 5065–5072; doi: 10.1242/dev.00774

Camp, D., Haitian He, HB., Li, S., Althaus, IW., Holtz, AM., Allen, BL., Charron, F., van Meyel, DJ. (2014). Ihog and Boi elicit Hh signaling via Ptc but do not aid Ptc in sequestering the Hh ligand. Development 141(20): 3879--3888.

Chial, H. (2008). Somatic mosaicism and chromosomal disorders. Nature Education 1(1):69

Clevers, H. (2006). Wnt/beta-catenin signaling in development and disease. Cell 127(3): 469–480.

Cook, RK., Christensen, SJ., Deal, JA., Coburn, RA., Deal, ME., Gresens, JM., Kaufman, TC., Cook, KR. (2012). The generation of chromosomal deletions to provide extensive coverage and subdivision of the Drosophila melanogaster genome. Genome Biol. 13(3): R21.

Donohoe CD, Csordás G, Correia A, Jindra M, Klein C, Habermann B, et al. (2018). Atf3 links loss of epithelial polarity to defects in cell differentiation and cytoarchitecture. PLoS Genet.14 (3): e1007241. https://doi.org/10.1371/journal. pgen.1007241.

Frappaolo, A, Sechi, S, Kumagai, T, Karimpour-Ghahnavieh, A, Tiemeyer, M, Giansanti, MG. (2018). Modeling Congential Disorders of N-Linked Glycoprotein Glycosylation in Drosophila melanogaster. Front. Genet. 9():436.

Franco, B., Bogdanik, L., Bobinnec, Y., Debec, A., Bockaert, J., Parmentier, M.L., Grau, Y. (2004). Shaggy, the homolog of glycogen synthase kinase 3, controls neuromuscular junction growth in Drosophila. J. Neurosci. 24(29): 6573--6577.

Fortunato, EA and Spector DH (2003). Viral induction of site- specific chromosome damage. Rev Med Virol. 13(1) pg21–37. https://doi.org/10.1002/rmv.368.

Germani, F., Bergantinos C and Johnston LA(2018). Mosaic Analysis in Drosophila Genetics 208(2) 473–490 ;https://doi.org/10.1534/genetics.117.300256.

Giansanti MG and Fuller MT (2012). What Drosophila spermatocytes tell us about the mechanisms underlying cytokinesis. Cytoskeleton (Hoboken). 69(11):869– 81. doi: 10.1002/cm.21063.

Gibson, MC. and Perrimon N (2005). Extrusion and Death of DPP/BMP- Compromised Epithelial Cells in the Developing Drosophila Wing. Science, 307(5716), 1785–1789. doi:10.1126/science.1104751

Gumbiner BM. (1992). Epithelial morphogenesis. Cell. 1;69(3):385–7.

Hsia, EYC., Zhang, Y., Tran, HS., Lim, A., Chou, YH., Lan, G., Beachy, PA., Zheng, X. (2017). Hedgehog mediated degradation of Ihog adhesion proteins modulates cell segregation in Drosophila wing imaginal discs. Nat. Commun. 8(1): 1275.

Knust E (2005). Development of Epithelial Cell Polarity in Drosophila Volume 7 Chapter 2 p121. Edited by (Birchmeier W and Birchmeier C, Epithelial Morphogenesis In Development And Disease(Cell adhesion and communication) ISBN 0-203-30376-8

Lee, T and Luo L. (1999). Mosaic analysis with a repressible cell marker for studies of gene function in neuronal morphogenesis. Neuron. 22(3):451–61.

Lee, T, Luo L. (2001). Mosaic analysis with a repressible cell marker (MARCM) for Drosophila neural development. Trends Neurosci. 24(5):251–4.

Olson, MV (1999). When less is more: Gene loss as an engine of evolutionary change. Am J Hum Genet 64: 18–23.

Olson, MV and Varki A (2003) Sequencing the chimpanzee genome: Insights into human evolution and disease. Nat Rev Genet 4: 20–28.

Paciorkowski, AR, Keppler-Noreuil K, Robinson L, et al. (2013). Deletion 16p13.11 uncovers NDE1 mutations on the non-deleted homolog and extends the spectrum of severe microcephaly to include fetal brain disruption. Am J Med Genet A. 161A(7):1523–1530.

Parks, AL., Cook, KR., Belvin, M., Dompe, NA., Fawcett, R., Huppert, K., etal., (2004). Systematic generation of high-resolution deletion coverage of the Drosophila melanogaster genome. Nat Genet. 36(3):288–92.

Paschinger, K., Straudacher, E., Stemmer, U., Fabini, G., Wilson, I.B.H. (2005). Fucosyltransferase substrate specificity and the order of fucosylation in invertebrates. Glycobiology 15(5): 463--474.

Price, MA and Kalderon, D. (2002). Proteolysis of the Hedgehog signaling effector Cubitus interruptus requires phosphorylation by Glycogen Synthase Kinase 3 and Casein Kinase 1. Cell 108(6): 823--835.

Powell, K and Piddini, E (2016). Chasing how cells out compete one another. Journal of Cell Biology. 213(3): 291–292.

Rauch, A; Schellmoser, S; Kraus, C; Dörr, HG; Trautmann, U; Altherr, MR; Pfeiffer, RA; Reis, A (2001). First known microdeletion within the Wolf-Hirschhorn syndrome critical region refines genotype-phenotype correlation. American Journal of Medical Genetics. 99(4): 338–42. doi:10.1002/ajmg.1203. PMID 11252005.

Reiter, LT., Potocki, L., Chien, S., Gribskov, M., Bier E. (2001). A systematic analysis of human disease-associated gene sequences in Drosophila melanogaster. Genome Res. 11(6):1114–25.

Roos, C., Kolmer, M., Mattila, P., Renkonen, R. (2002). Composition of Drosophila melanogaster proteome involved in fucosylated glycan metabolism. J. Biol. Chem. 277(5): 3168--3175.

Rodríguez, AV, Didiano D, Desplan C (2012). Power tools for gene expression and clonal analysis in Drosophila Nature Methods 9: 47–55.

Roote, J and Russell, S. (2012). Toward a complete Drosophila deficiency kit. Genome Biol. 13(3): 149.

Ruddle, FH, Bentleyl, KL, Murtha MT, Risch N (1994). Gene loss and gain in the evolution of the vertebrates. Development Supp. 155-161.

Ryder, E., Blows, F., Ashburner, M., Bautista-Llacer, R., Coulson, D., Drummond, J., Webster, J., et al., (2004). The DrosDel collection: a set of P-element insertions for generating custom chromosomal aberrations in Drosophila melanogaster. Genetics 167(2): 797–813.

Ryder, E., Ashburner, M., Bautista-Llacer, R., Drummond, J., Webster, J., Johnson, G., et al., (2007). The DrosDel deletion collection: a Drosophila genomewide chromosomal deficiency resource. Genetics. 177(1):615–29.

Schock, F and Perrimon N. (2002). Molecular mechanisms of epithelial morphogenesis. Annu Rev Cell Dev Biol. 18:463–93. DOI: 10.1146/annurev.cellbio.18.022602.131838.

Shen, J and Dahmann C (2005). Extrusion of Cells with Inappropriate Dpp Signaling from Drosophila Wing Disc Epithelia. Science, 307(5716), 1789–1790. doi:10.1126/science.1104784.

Smith-Bolton, RK., Worley, MI., Kanda, H and Hariharan, IK. (2009) Regenerative growth in Drosophila imaginal discs is regulated by Wingless and Myc. Dev. Cell 16: 797–809.

Slavotinek, A. (2012). Microdeletion Syndromes. In eLS, (Ed.). doi:10.1002/9780470015902.a0005549.pub2

St Johnston, D., (2002). The art and design of genetic screens: Drosophila melanogaster. Nat. Rev. Genet. 3: 176–188.

Sustar A., Bonvin M., Schubiger M and Schubiger G (2011) Drosophila twin spot clones reveal cell division dynamics in regenerating imaginal discs. Developmental Biology (356) 2: 576–587.

Szuperak M, Salah S, Meyer E, Usha N, Ikmi A and Gibson M. (2011) Feedback modulation of Drosophila BMP signaling by the novel extracellular protein, Larval Translucida Development 138, 715–724.

Nagarajan U, Pakkiriswami S and Pillai AB. (2015) Sugar tags and tumorigenesis. Frontiers in Cell and Developmental Biology 3(69), 1–5.

Usha N and Shashidhara LS. (2010) Interaction between Ataxin-2 Binding Protein 1 and Cubitus- interruptus during wing development in Drosophila. Developmental Biology 341, 389–399.

Yamamoto S, Jaiswal M, Charng WL, Gambin T, Karaca E, Mirzaa G, et al., (2014). A drosophila genetic resource of mutants to study mechanisms underlying human genetic diseases. Cell. 25;159(1):200–214. doi: 10.1016/j.cell.2014.09.002.

Wu, JS and Luo L. (2006). A protocol for mosaic analysis with a repressible cell marker (MARCM) in Drosophila. Nat Protoc. 1(6):2583–9.

Zhang W, Hong M, Bae GU, Kang JS, Krauss RS. (2011) Boc modifies the holoprosencephaly spectrum of Cdo mutant mice. Dis Model Mech. 4(3):368–80. doi: 10.1242/dmm.005744.

